# Priority effects determine community composition at the strain level in the honeybee gut microbiota

**DOI:** 10.1101/2025.11.22.689826

**Authors:** Aiswarya Prasad, Gonçalo Santos-Matos, Alexandra Szigeti-Genoud, Florent Mazel, Philipp Engel

## Abstract

Gut microbial communities often differ at the strain level among individual hosts, but the mechanisms driving this variation remain poorly understood. One potential factor is priority effects, a process in which differences in the timing and order of microbial colonization influence subsequent community assembly (“first come, first served” dynamics). We hypothesize that such priority effects operate at the strain level within species, where closely related bacteria exhibit niche overlap, and that these dynamics can lead to community divergence even under similar environmental conditions. We tested these predictions, using the gut microbiota of honeybees, which harbor conserved microbial communities that differ in strain composition among individual bees. We sequentially colonized microbiota-depleted honeybees with two distinct microbial communities composed of the same twelve core microbiota species but different strains, ensuring that individuals shared species-level composition but differed at the strain level. We found that firstcomer strains consistently dominated the resulting communities, suggesting strong priority effects. Dropout experiments in which the firstcomer strain of a species was removed led to only partial increases in the colonization success of the conspecific latecomer, suggesting that both intra- and inter-species interactions contribute to priority effects. Our findings highlight the significant role of priority effects in strain-level community assembly and reveal their influence in shaping the specialized gut microbiota of honeybees, with important implications for the development of probiotic strategies in beekeeping.

**GRAPHICAL ABSTRACT:** 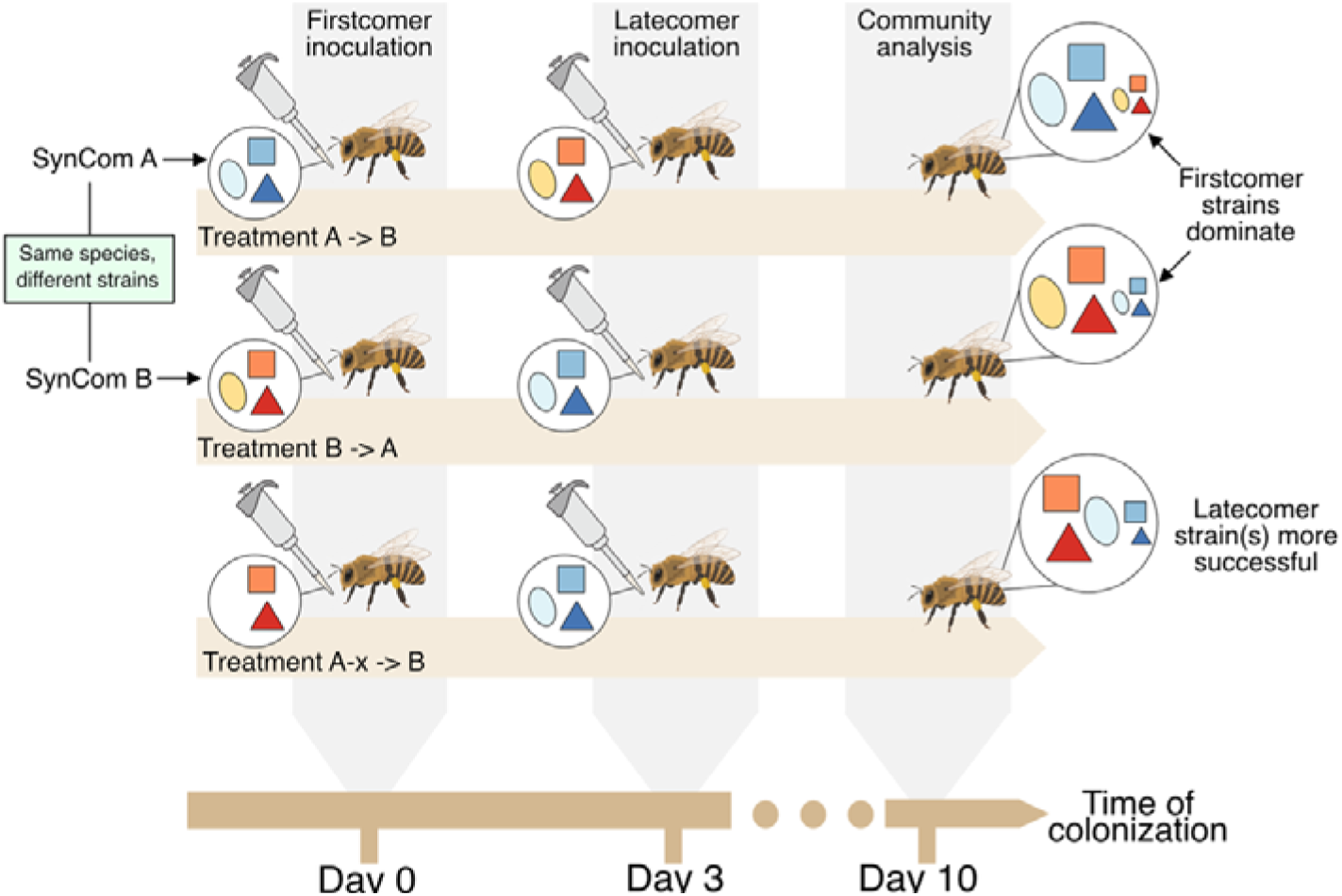

## INTRODUCTION

Gut microbial communities exhibit substantial bacterial diversity, particularly at the strain level (Ellegaard & Engel, 2019; Ley et al., 2008; Schloissnig et al., 2013). Closely related strains can stably co-exist within the same host (Ellegaard & Engel, 2019; Garud et al., 2019; Truong et al., 2017; Wolff et al., 2023). But different hosts typically harbour distinct strain profiles, even when closely related or exposed to the same environment (Garud et al., 2019; Wolff et al., 2023). These differences in strain-level composition can have significant implications for the functional potential of the microbiota and their interactions with and benefits to the host (Yan et al., 2020). Studying the mechanisms that underscore strain-level differences across individuals are hence fundamental to our understanding of the relationship between hosts and their gut microbiota.

An important mechanism of microbial community assembly is environmental filtering. This occurs when differences in physicochemical conditions, such as nutrient availability, pH, transit time, oxygen levels, and immune system activity determine which microbes can colonize a given environment based on their functional traits. These deterministic factors have been relatively well studied in the gut microbiota (Mallott & Amato, 2021; Miller & Bäumler, 2021; D. D. Sprockett et al., 2023), but other mechanisms are less understood. These include dispersal limitation, or the ability of microbes to reach and colonize specific hosts (due to spatial barriers such as geography) (Mazel et al., 2023; Moeller et al., 2017), and stochastic processes, random colonization events that occur independently of microbial traits or environmental conditions (Nemergut et al., 2013; Zhou & Ning, 2017). One such stochastic phenomenon is priority effects (Debray et al., 2022), where the timing and order of arrival shape the resulting community composition (Gleason, 1927; Nemergut et al., 2013). Microbes that arrive earlier can occupy and/or alter the environment, affecting the colonization success of subsequent arrivals. While microbiome studies have investigated priority effects between bacterial species (Carlström et al., 2019; Martínez et al., 2018; Ojima et al., 2022; D. Sprockett et al., 2018), few studies have documented priority effects at the strain level and in the context of a multi-species community (Chen et al., 2024; Segura Munoz et al., 2022). This is surprising given that contemporary coexistence theory (Adler et al., 2007; Chesson, 2000; Fukami et al., 2016; Mayfield & Levine, 2010; Orr et al., 2025) predicts priority effects should be stronger between competitors with a small fitness differences and overlapping niches. The presence of a complex, multi-species community can further amplify these effects, as interactions with other community members may impose additional constraints on the niches of each strain in the community. Hence, we hypothesize that priority effects should be an important factor in shaping gut microbiome composition at the strain level leading to differences across individuals.

Western honeybees (*Apis mellifera*) are an excellent model for studying the processes underlying community assembly, particularly priority effects. Honeybees harbor a relatively simple and stable gut microbiota, composed of a few host-specific bacteria that include multiple closely related species and strains (Ellegaard & Engel, 2019; Engel et al., 2012; Kwong & Moran, 2016). While most species are consistently present and often co-occur in individual bees, strain-level variation is more individualized, i.e. conspecific strains (i.e. strains of the same species) tend to segregate between individual bees, even nestmates from the same colony (Ellegaard & Engel, 2019). Priority effects early in life, when newly emerged adult bees acquire the microbiota from their nestmates, have been proposed as one possible mechanism for the observed patterns (Ellegaard & Engel, 2019).

In this study, we experimentally tested the role of priority effects in gut community assembly at the strain level by inoculating microbiota-deprived (MD) bees under laboratory conditions with two microbial communities administered three days apart. Each community consisted of the same twelve species, but different strains. We also conducted dropout experiments, in which a single strain from a subset of the 12 species was excluded from the firstcomer community, to test whether this increased the colonization success of its conspecific strain in the latecomer community. Our results show that priority effects are an important ecological process shaping gut microbiota diversity at the strain level and that they are influenced by both within- and between species interactions.

## MATERIALS AND METHODS

### EXPERIMENTAL DESIGN

To test the role of priority effects in shaping strain-level community assembly in the honeybee gut microbiota, we designed an experiment where strains from the same species were alternatively introduced as “firstcomers” or “latecomers”, three days apart. If priority effects were strong, we expected the firstcomer to predominate in the community. We assembled multi-species synthetic communities comprised of the same 12 species but harbouring a different strain for each species. All species belonged to either *Bifidobacterium*, *Lactobacillus,* or *Bombilactobacillus* which are the predominant genera in the distal hindgut (rectum) of the Western honeybee (*Apis mellifera*) (**Fig. 1A, Supplementary Table S1**). MD bees were inoculated at Day 0 with “community A” (firstcomer strains) and then challenged with “community B” (latecomer strains) at Day 3 (**Fig. 1D**). The timing of the second inoculation was chosen to be early enough that the community has not yet been fully established. We validated in a pilot experiment, that inoculation at Day 3 resulted in successful colonization meaning that bacteria do not colonize less successfully just because they arrive later when no previous community was inoculated (see **Supplementary methods** for details). The priority effect was measured for each strain in a given community as the ratio of its absolute abundance when inoculated within the context of the community on Day 0 and on Day 3 (**Fig. 1B**). To replicate the measurement of priority effect at the strain level but under different community backgrounds, we made six community combinations by shuffling strains between community A and B such that in each combination, a given strain co-occurred with a different set of strains of the other 11 species. (**Supplementary Table S2**, **Fig. 1C**). For two *Bombilactobacillus* species (*B. mellifer*, *B. mellis*) and one Bifidobacterium species (*B. cornyforme*), the same strain was used in all communities, and for one Lactobacillus species (*L. melliventris*) we had by mistake included two strains in some of the communities (**see Supplementary methods** for a more detailed explanation). So, we could only assess the priority effects for 8 of the 12 tested species in a reciprocal manner.

**Figure 1.**
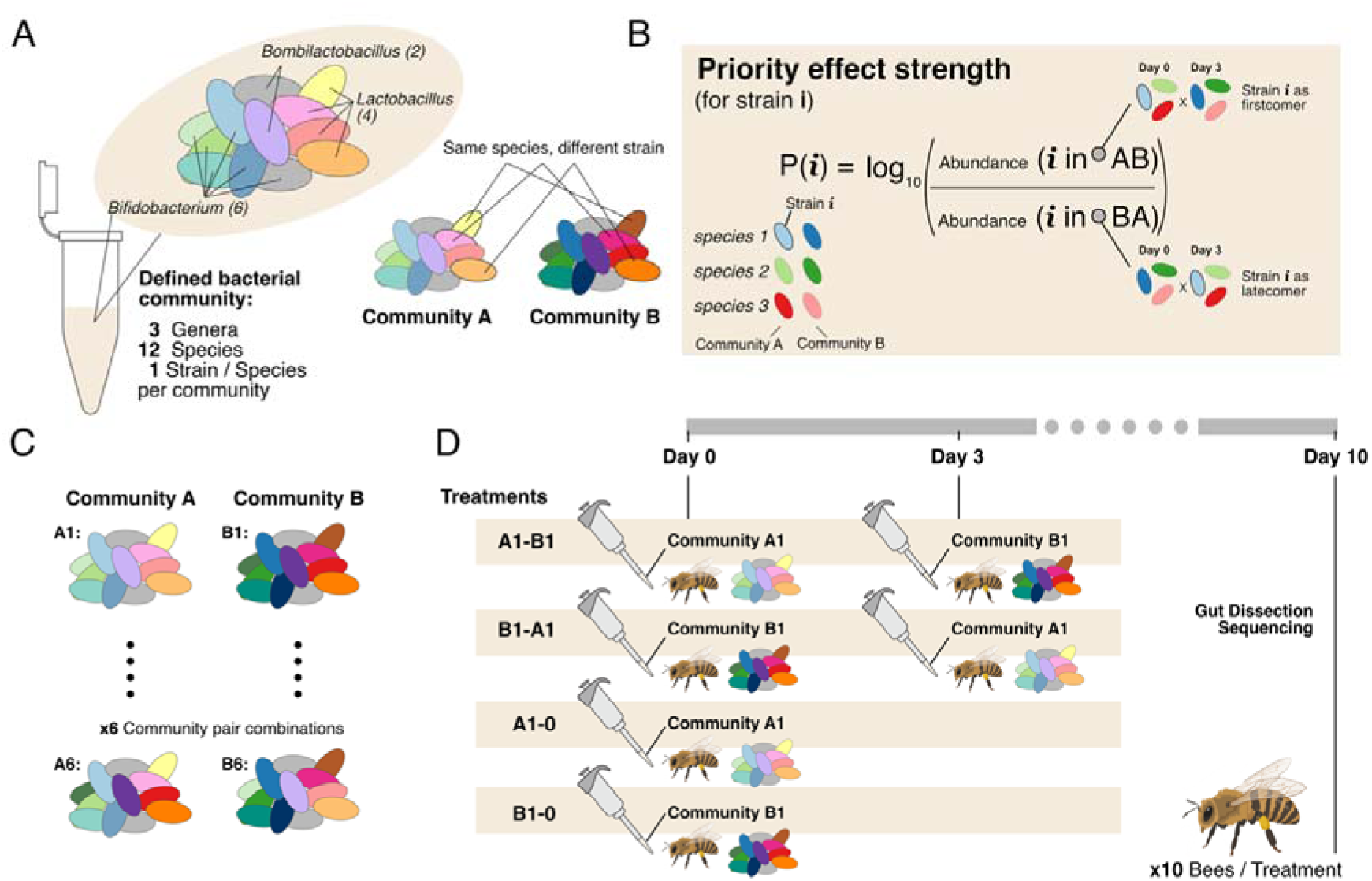
Experimental design and community selection. (A) Schematic of the multi-species bacterial communities used for the experiments. Each community consisted of twelve species, two of Bombilactobacillus, four of Lactobacillus, and six of Bifidobacterium. (B) Illustration of the approach used to measure the priority effect for a given strain *i*. (C) Schematic representing community pairs A and B made in 6 different combinations (1 – 6) by shuffling strains between them. (D) Illustration of the experimental design to test each community combination (here, 1).

For each pair of community combinations, we included four treatments (A, B, AB and BA) (**Fig. 1D**). In treatments A and B, MD bees were only inoculated with one community at Day 0. These treatments serve as controls to determine community composition and the abundance of each strain when inoculated without a secondary challenge on Day 3. In AB and BA, the two communities were inoculated one after the other on Day 0 and Day 3, respectively (**Fig. 1D**). Each strain appeared in either treatment A or B of each community combination. Consequently, a strain present in community A, is a firstcomer in treatment AB and a latecomer in treatment BA. For each community combination and treatment, we analyzed 8-10 bees from the cage. This way, each strain was measured a total of about 60 times as a firstcomer and a latecomer each across the six different community combinations (**Fig. 1C**).

### BACTERIAL CULTURING AND SYNTHETIC COMMUNITY ASSEMBLY

A total of twenty-two strains belonging to the twelve selected species were used in this study. They were all isolated from the gut of adult bees of the Western honeybee (*Apis mellifera*) either in this or in previous studies (Brochet et al., 2021; Ellegaard et al., 2019). We used pairwise average nucleotide to determine which strain belongs to which species (**Supplementary Fig. S1**). Strains for which no complete genome was available were resequenced using PacBio or Nanopore sequencing and all full length16S rRNA of each strain were extracted using barrnap to ensure that all strains can be discriminated by at least 1-2 SNPs in at least one variant. Details about each strain can be found in **Supplementary Table S1**. To assembly the six synthetic communities, the selected strains were grown on solid De Man – Rogosa – Sharpe agar (MRSA) (supplemented with 2% w/v fructose and 0.2% w/v L-cysteine-HCl) in petri dishes at 34°C under anaerobic conditions. After ∼48h of growth, each strain was harvested using a sterile loop and resuspended in 300µL of a sterile solution of PBS+20% glycerol. The solution was diluted to an OD_600_ value of 2 in PBS+20% glycerol. To assemble the synthetic communities, 10 µL of each of the twelve strains were mixed into a sterile tube. This resulted in about 120µL for each community combination (for dropout communities, sterile PBS+20% glycerol solution was added instead of the dropped strain), to which 80 µL of PBS+20% glycerol solution were added to make up a total of 200 µL of microbial solution for each community. This process ensured that each strain would be present in a quantity equivalent to a final OD_600_ value of 0.1. The final mixture for each community was thoroughly mixed and separated into 50µL aliquots, which were flash frozen using liquid nitrogen and stored at −80°C until used.

### GNOTOBIOTIC BEE EXPERIMENTS

Microbiota-deprived (MD) honeybees were obtained from *Apis mellifera carnica* colonies maintained at the University of Lausanne, as previously described (Kešnerová et al., 2017). One day after eclosion (here D0), bees were fed 5 µL of inoculum prepared from glycerol stock aliquots of the synthetic community, diluted in 450 µL of a 1:1 mixture of sterile sugar water (50% sucrose, w/v) and 1× PBS. All inoculations were performed at room temperature. After inoculation, bees were transferred to their respective sterilized 3D-printed cages (one per treatment), containing sterile pollen and sugar water tubes *ad libitum*. In each treatment, 10 – 12 bees were included per cage. For treatments involving a second inoculation on Day 3, feeding was repeated as done on Day 0. Finally, on Day 10, all surviving honeybees (8 – 10) were sacrificed and their hindguts were dissected, flash frozen using liquid nitrogen, and stored at −80°C until DNA extraction.

### DNA EXTRACTION AND AMPLICON SEQUENCING

DNA was extracted from dissected bee guts using a magnetic bead-based protocol optimized for high-throughput processing on the Opentrons OT-2 liquid-handling robot. Samples were homogenized in bead-beating tubes containing a mix of 1□mm glass beads and 0.1□mm zirconia beads, 750 μL 1× G2 buffer supplemented with lysozyme (100□mg/mL). For lysis the homogenate was then incubated at 37□°C for 30□min with shaking (900□rpm), followed by the addition of Proteinase K (20□mg/mL) and further incubation at 56□°C for 1□h. For extracting DNA from the inocula fed to bees, 165 μL was used for beadbeating. Lysates were centrifuged, and 80□μL was transferred to a 96-well PCR plate. Extraction blanks were included as wells only containing nuclease-free water instead of gut homogenates. DNA purification was performed using CleanNGS magnetic beads (Clean NA #CNGS-0050) on an Opentrons® OT-2 robot. Automated steps included magnetic bead binding, two 80% ethanol washes, drying, and elution in nuclease-free water. Eluates (30□μL) were collected in a new plate and stored at −20□°C or −80□°C. DNA concentrations were quantified using a Qubit™ fluorometer and the 1X dsDNA HS assay. 0.1x dilutions were made and stored in an additional plate and used for further steps.

### QUANTIFICATION OF TOTAL BACTERIAL ABUNDANCE

qPCR was performed on DNA extracts diluted to 0.1x to minimize the effect of contaminants. Primers targeting a 162 bp stretch of the V4 region of the 16S rRNA gene were used to quantify the entire bacterial community as described before (Kešnerová et al., 2017), and as outlined in the **Supplementary Methods**.

### LIBRARY PREPARATION AND AMPLICON SEQUENCING

For amplicon sequencing, multiplexed amplicon libraries were prepared using the “Amplification of Full-Length 16S Gene with Barcoded Primers for Multiplexed SMRTbell® Library Preparation and Sequencing protocol” (Version 05). PCR amplification was performed using the KAPA HiFi HotStart ReadyMix (KAPA Biosystems) and a pre-prepared barcoded primer plate containing the forward and reverse primers 5’-GCATC/barcode/AGRGTTYGATYMTGGCTCAG-3’ and 5’-GCATC/barcode/RGYTACCTTGTTACGACTT-3’ in each well combined to result 96 unique pairs as recommended in the protocol. Amplicons were diluted to 0.01x and quantified using qPCR as described in the following section and the **Supplementary Methods**. Using the results of the quantification, amplicons were pooled in equimolar quantities for library preparation using the SMRTbell Prep Kit 3.0. Library preparation and sequencing were carried out in the Next Generation Sequencing Platform of the University of Bern.

### AMPLICON SEQUENCING ANALYSIS TO DETERMINE STRAIN ABUNDANCE

Raw reads from sequencing were processed using DADA2 v1.30.0 (Callahan et al., 2016) to infer amplicon sequence variants (ASV) and their counts per sample as described before (Callahan et al., 2019). The detailed code used for this analysis is provided in the associated GitHub repository https://github.com/Aiswarya-prasad/2025_aprasad_PriorityEffects. Importantly, the *removePrimers* function was used for primer removal and automatically re-orienting all reads in the forward direction. Reads were then filtered to only keep reads in the range of 1000 – 1600 bp. Finally reads were dereplicated and denoised using a PacBio sequencing specific error model.

The inferred ASVs were matched with unique 16S rRNA variants of each strain and used to infer the counts per strain that would result in the observed ASV counts table generated by DADA2. This was done using the code (infer_species_counts.py) provided in the associated GitHub repository https://github.com/Aiswarya-prasad/2025_aprasad_PriorityEffects. To resolve strain-level composition from ASV counts, we implemented a matrix-based least-squares approach, leveraging known relationships between strains and their 16S rRNA gene copy profiles. We took this approach because each strain comprises several unique or non-unique ASVs and the counts of their ASVs are not equal even when they have the same copy number due to technical errors in amplification and sequencing (see **Supplementary Methods** for more details).

Samples with low 16S rRNA gene copies (C_T_> 26.53 or undetermined) or sequencing depth (<100 known reads) were removed from further analyses (n = 15 of 365). The relative abundance of each strain was calculated as the ratio of number of reads assigned to that strain and the total number of reads in that sample. The absolute abundance of each strain in a sample was estimated as the product of relative abundance and total copy number for that sample. The detection limit of each sample was the number of copies yielding 1 read in that sample, *i.e.*, the ratio of total 16S copies and the number of reads sequenced. Any values of abundance below this were considered below the limit and their value was set to 1 (for plotting on a log scale without errors).

### STATISTICAL ANALYSES

To compare community profiles across treatments, PERMANOVA was used. To do this, we used Vegan (v2.6-4) package to obtain Bray-Curtis distances from the log-transformed sample vs. bacterial abundance matrix. The distance matrix was then used to make a PCoA and the adonis2 function used to perform the PERMANOVA analysis. The effect size (Ω^2^) was calculated using the adonis_OmegaSq function from the package https://github.com/Russel88/MicEco. The results are denoted as Ω^2^ and p-values.

The Wilcox rank sum test in R was used to compare differences between treatments as stated in the respective figure descriptions or references. The Benjamini-Hochberg method was used to correct for multiple comparisons where necessary. All tests were two-tailed and a p-value < 0.05 was considered statistically significant.

For the generalized linear mixed model (GLMM), we can use the *glmer* function from the lme4 package in R. The model was set up to predict the transformed absolute counts based on the arrival order, while accounting for random effects due to different community combinations. It was carried using It was carried out using a Gamma distribution with a log link function, which is appropriate for modelling positive continuous data like counts.

## Results

### PRIORITY EFFECTS DETERMINE COMMUNITY COMPOSITION AT THE STRAIN LEVEL

Most honeybees across treatments were successfully colonized by the inoculated community, (see total bacterial abundances in **Supplementary Fig. S2**). The abundance of bacteria in the gut of honeybees at the sampling day (i.e. Day 10) was orders of magnitude higher than in the inoculum (fed on Day 0 and Day 3) and in non-inoculated MD bees fed sterile sugar water and pollen, indicating that the strains in the inoculum proliferated within the honeybee gut resulting in the assembled communities (**Supplementary Fig. S3**).

All twelve species were detected in most honeybee samples and the colonizing strains matched those present in the inocula of each treatment, confirming both the success of the experimental setup and the gnotobiotic status of our bees (**Supplementary Fig. S4 and S5**). Total bacterial loads in each sample were not significantly correlated with the number of strains present (Spearman correlation ρ = 0.02, p = 0.219) (**Supplementary Fig. S6**), and the summed abundance for each species did not differ significantly whether one or two strains of the species were inoculated in the treatment (**Supplementary Fig. S7**). This indicates that the strains in each community reach carrying capacity and that latecomer strains compete in the niches occupied by the firstcomers rather than occupying additional empty niches.

To determine whether priority effects shape overall community composition (**Fig. 2**), we used the abundance of strains to estimate the Bray-Curtis compositional distance between the samples of the four treatments in each community combination (∼10 bees per treatment). PERMANOVA tests confirmed a strong (Ω^2^ > 0.8) and significant (*p* < 0.05) effect of treatment across all community combinations. Communities AB (A inoculated at day 0 and B at day 3) closely resembled communities A, while communities BA resembled communities B across all replicates (**Fig. 2**). These differences were also visible across all replicates by metrics using either relative abundances or absence/presence of strains to compare the communities (**Supplementary Fig. S8**). These results indicate that the firstcomer strains dominated the communities and that the order of arrival of strains determines the overall community composition (**Fig. 2**).

**Figure 2.**
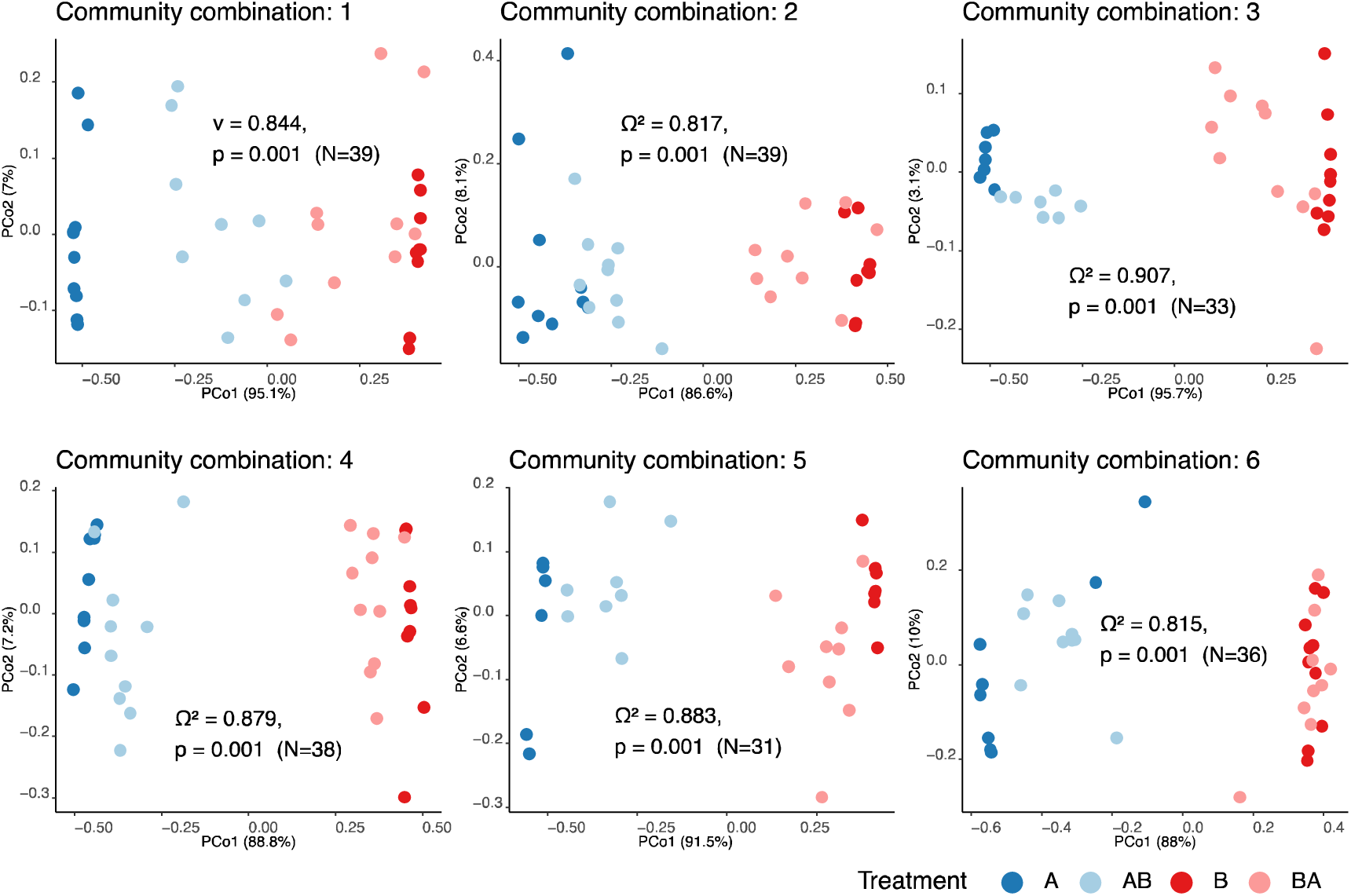
Community composition across samples. PCoA plots to visualize Bray-Curtis distance estimated from the matrix of log-transformed absolute abundance of bacterial strains across community combinations, with color indicating the treatment (A and B, community A and B on Day 0, respectively; AB, community A on Day 0 and community B on Day 3, and vice versa for BA).

### PRIORITY EFFECTS ARE CONSISTENTLY OBSERVED FOR ALL STRAINS, BUT VARY IN STRENGTH

Strains of the same species may vary in the extent to which their niches overlap or in their ability to interfere with already established strains which can influence the strength of priority effects (Debray et al., 2022; Fukami et al., 2016; Peay et al., 2011). Therefore, we next aimed to assess whether there are differences in the magnitude of the priority effects on individual strains within the tested communities. All strains colonized at significantly lower abundances (Wilcox rank sum test, p < 0.05) when they were inoculated on Day 3 as latecomers (A in BA, or B in AB) than when inoculated on Day 0 as firstcomers (i.e. A in AB, or B in BA) (**Fig. 3**) suggesting that priority effects act on all tested strains. Notably, several strains were nearly undetectable when introduced as latecomers (e.g., *Bifidobacterium asteroides* ESL017/ESL0822 and *Bifidobacterium* sp2 ESL0200/ESL0819; Fig. 3, Supplementary Fig. S9), while others, in particularly *Lactobacillus apis* ESL0263, were only marginally affected. A generalized linear mixed model confirmed that arriving second significantly reduced absolute abundance, relative to arriving first or being alone (p < 0.05 for second and p > 0.05 for first) for all strains except *Lactobacillus apis* ESL0263 (**Supplementary Table S5**). To quantify differences among strains, we calculated the priority effect strength in each community combination for a given strain as the log ratio of the median abundance of the strain when it was a firstcomer vs latecomer (**Fig. 1B, Supplementary Table S4**). As expected, priority effect strength was higher than zero for all strains (**Fig. 4**), but varied across strains, even among those of the same species (**Fig. 4**). Moreover, some strains showed substantial variation across replicates, while others did not, suggesting that in some cases the community background, and hence the interactions with allospecific strains, influenced priority effect strength.

**Figure 3.**
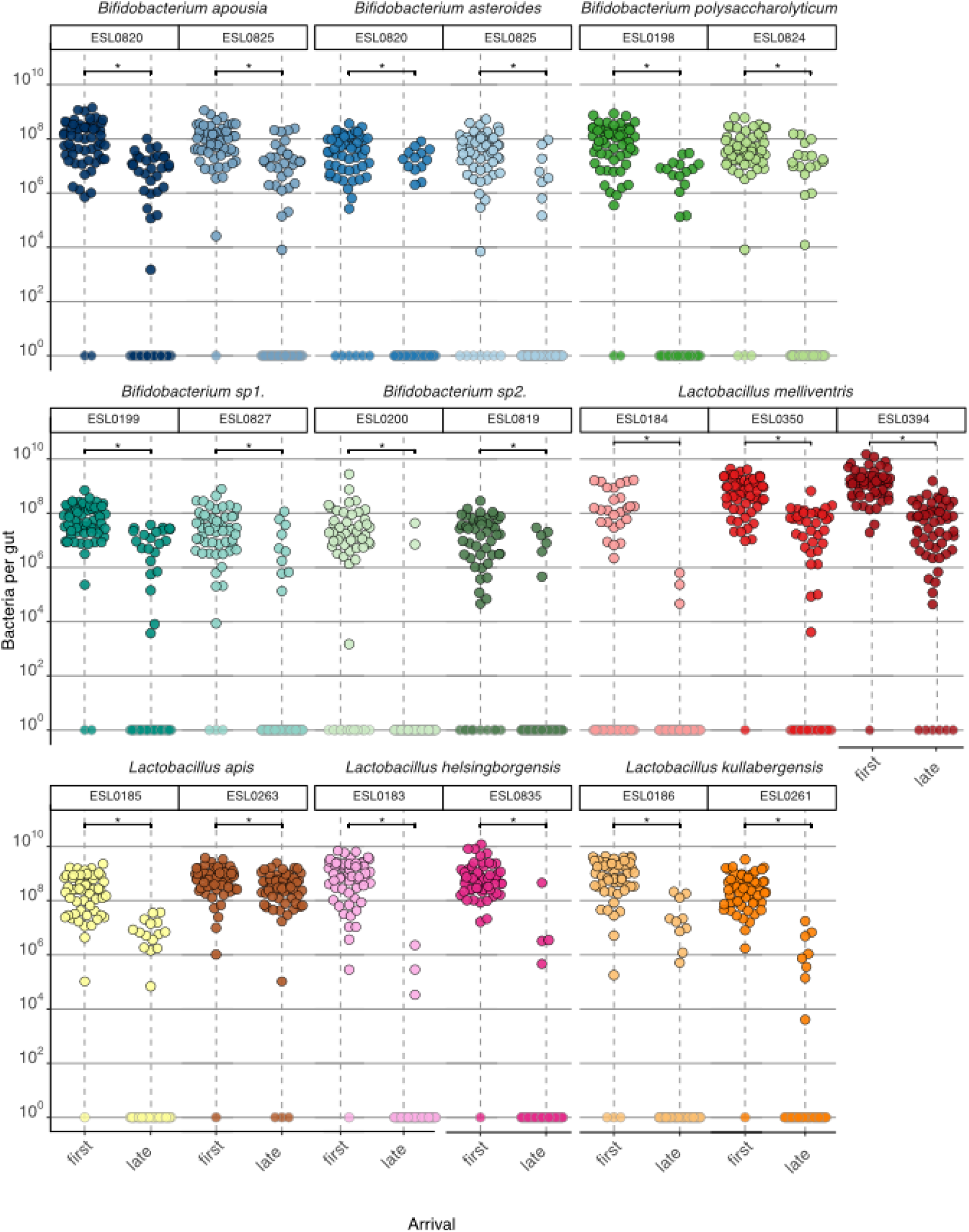
Abundance of each strain by arrival order. Absolute abundance of each strain (faceted by strain) under different arrival orders. “First” treatment: the focal strain was inoculated on Day□0 as part of a defined microbial community, and on Day□3 a second community (also containing the same conspecific strain) was added. “Late” treatment: the focal strain was instead introduced on Day□3 as part of the second community, into gnotobiotic bees that had already been inoculated on Day□0 with a community containing the same conspecific strain. Each plot includes all the treatments across the six community combinations. Result of Wilcoxon rank sum test (two-sided) are annotated (* - p<0.05), complete results of statistical test, including sample size per, are included separately (Supplementary Table S3). Points with grey outlines represent samples where the strain was not detected (below the detection threshold for that sample) and hence set to a value of 1.

**Figure 4.**
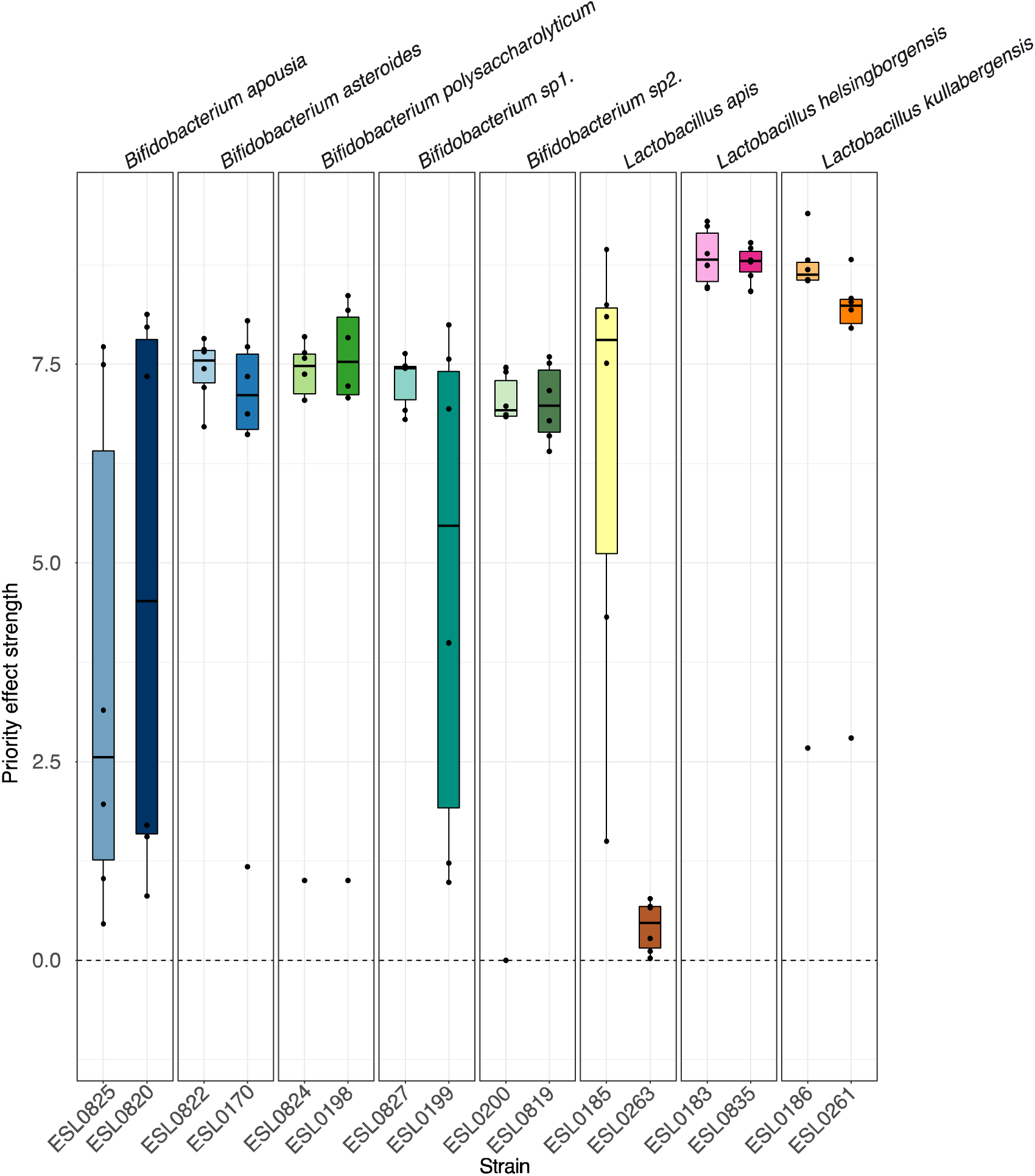
Strength of priority effect by strain across community combinations. Each point (black dot) represents a particular strain tested within one of the six community-pair combinations. The strength of the priority effect is computed as shown in the formula in Fig. 1B using the median absolute abundance of a given strain in ∼10 bees of each treatment AB and BA. Zero represents the value corresponding to no priority effect.

### Conspecifics account for only part of the observed priority effects

We hypothesized that priority effects are primarily mediated by within-species (conspecific) competition, rather than between-species (allospecific) interactions, based on the assumption that conspecific strains exhibit greater niche overlap. To test this, we used the same experimental setup as before but modified the firstcomer community by omitting the strain of one focal species. At Day 3, we introduced the full latecomer community (**Fig. 5A**). This experiment was performed using three randomly selected species from the original set of 12 (*Bifidobacterium apousia*, Bifidobacterium sp1., and *Lactobacillus apis*), for which both strains were removed in reciprocal order (i.e., dropping out the firstcomer strain in the AB and BA treatment, respectively). To assess how the absence of the firstcomer strain affected colonization of the conspecific latecomer strain, we compared its absolute abundance in the dropout treatment to its abundance when introduced alongside the full community (either as firstcomer or latecomer). We expected that removing the firstcomer strain would increase colonization success of the latecomer conspecific strain, compared to the condition in which the full firstcomer community was present (**Fig. 5A**).

**Figure 5.**
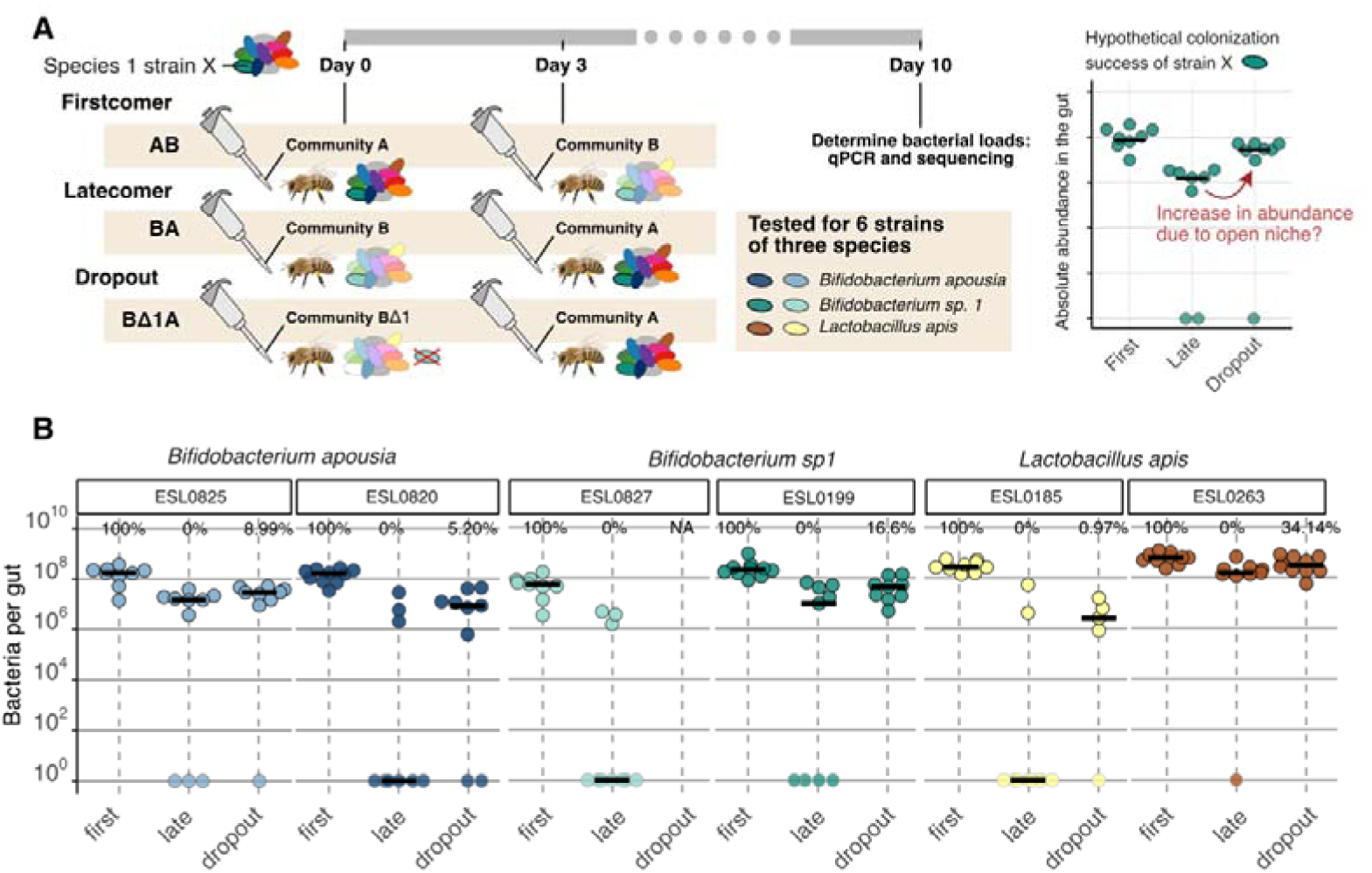
Colonization success of strains as latecomers in full and dropout treatments. (A) Scheme of the experimental design (left panel) and theoretical example of the the analysis (right panel). (B) Absolute abundance of six tested strains of three species when arriving first (‘first’) or late (‘late’) in the full community versus when arriving late in the dropout treatment (‘dropout’) in which the conspecific strain was dropped out from the firstcomer community. The percentages indicate the median colonization level of the focal strain relative to the median colonization level in the firstcomer community (100%) and the latecomer community (0%) using the formula: (abundance(dropout) – abundance(late)) / (abundance(first) – abundance(late)). Samples in which the strain was above the detection limit were considered for this calculation. The effect of the conspecific dropout on the abundance of strain ESL0827 of *Bifidobacterium sp1.* could not be determined due to low sequencing depth (**Supplementary Fig. S2**)

All latecomer strains, except for ESL0185 of *Lactobacillus apis*, showed improved colonization success when bees were colonized with the dropout firstcomer community compared to the full firstcomer community (**Fig. 5B**, ‘late’ vs ‘dropout’). However, none of the latecomer strains reached the same abundance levels as when they were introduced as part of the firstcomer community (**Fig. 5B**, ‘first’): they reached between 1%-50% of the median abundance when introduced with the firstcomer community depending on the strain. Our sequencing analysis also revealed that latecomer strains of species other than the one dropped out were affected. In particular, several *Bifidobacterium* species in treatments where the dropouts were *Bifidobacterium* strains (**Fig. 5, Supplementary Table S6**), were more successful in colonization in those two treatments than in the full community or other dropout treatments (**Fig. 5**, **Supplementary Table S6**). Based on these results, we conclude that priority effects are influenced not only by strain interactions within species (i.e., conspecific interactions) but also by interactions between species, particularly among closely related species.

## DISCUSSION

Empirical evidences are beginning to show that the order of arrival (*i.e.*, priority effects) plays an important role in shaping community assembly in microbial ecosystems (Boyle et al., 2021; Carlström et al., 2019; Debray et al., 2023; Garrido-Sanz & Keel, 2024; Peay et al., 2011) including gut microbiomes (Chen et al., 2024; Gurung et al., 2024; Laursen & Roager, 2023; Ojima et al., 2022; Segura Munoz et al., 2022). However, few studies (e.g. Segura Munoz et al., 2022) have explored how priority effects influence the assembly of gut communities at the strain level and within the context of defined multi-species microbial consortia. By leveraging full-length 16S rRNA amplicon sequencing to distinguish between conspecific strains, we demonstrate that priority effects are key determinants of strain-level community assembly in the honeybee gut microbiota model.

We assessed the importance of priority effects by using a synthetic microbial community composed of several related species rather than individual strains (Großkopf & Soyer, 2014). This approach ensured that the observed effects were relevant in the presence of inter-species interactions, such as competition for space and nutrients, mimicking the natural environment in the gut. (Brochet et al., 2021; Ghoul & Mitri, 2016; Kešnerová et al., 2017). Further, our approach enabled us to simultaneously evaluate the impact of arrival order on several strains (two strains of eight species, i.e. a total of 16 strains) inoculated reciprocally as firstcomers and latecomers. To account for unexpected effects of interactions between allospecific strains in the same community, we tested several community combinations, i.e. the priority effect of each strain was assessed in the background of different allospecific strains.

Our findings demonstrate that priority effects play a significant role in shaping the strain level composition of the bee gut microbiota. When bees were sequentially inoculated with the two communities, their gut microbiota consistently resembled that of bees exposed only to the first community. This indicates that firstcomer strains effectively dominated the community, hindering the establishment of latercomer strains. Such priority effects may explain - or at least contribute to – the natural variation in strain-level composition observed among individual worker bees within a colony (Ellegaard & Engel, 2019). Similar early-life effects have been shown to drive microbiota variation in laboratory mouse models, even under tightly controlled host and environmental conditions (Martínez et al., 2018). Like in other animals, the gut microbiota of adult bees is acquired after ‘birth’ (i.e. pupal eclosion) through social interactions with nestmates. Therefore, the specific strains a newly emerged bee acquires likely depend on which nestmates it encounters during its early post-emergence period (firstcomer “seeding”). These initial interactions can thus result in persistent individual differences in microbiota composition and could even contribute to the host-specific nature of the bee gut microbiota (Mazel et al., 2025). This could be further tested in a natural setting by using tagged bacterial strains and tracking the interactions of newly emerged bees to compare the strain-level composition of their gut microbiota with that of the adults that they interacted with first as opposed to later in life.

Compared to other animals gut microbiomes, priority effects may be particularly strong in honeybees, as newly emerged adults spend the first few days inside the hive and only sporadically embark on defecation flights (Langstroth, 1853). As a result, food residence time in the gut may be relatively long, potentially leading to low strain-level turnover, and ecological opportunities for exposure to the microbiota of other honeybee species are low. However, as adult worker bees age, they transition from in-hive nurses to outside foragers, undergoing substantial physiological and behavioural changes. Importantly, during this shift they dramatically reduce pollen intake and increasingly rely on nectar and honey as their primary diet. Recent findings show that foragers harbour a distinct strain-level community compared to nurses (Baud et al., 2023). Therefore, even though early arriving strains might dominate the community in nurse bees, other processes and mechanisms than priority effects are likely at play, such as selection in the host gut environment or dietary differences, and may affect the community composition, *e.g.*, when bees transition to foragers.

Strains of the same species in the bee gut can differ substantially in their accessory gene content, leading to variation in functional potential (Ellegaard et al., 2019; Ellegaard & Engel, 2019; Engel et al., 2012; Van Rossum et al., 2020). As a result, they may influence the host in distinct ways – for example, through differences in carbohydrate degradation or the production of metabolites with neuroactive properties. Consequently, the order in which strains colonize the gut early in life, and the resulting differences in community composition at the strain-level, could have lasting effects on how the microbiota interacts with and impact the host. Such host effects of arrival order have been shown to occur in legume-rhizobium mutualism (Boyle et al., 2021) and may likely also play an important role in more complex microbial communities, such as those in the animal gut. Our results also suggest that probiotic strains, which are already used in bee management (Damico et al., 2025), need to be administered early in the bee’s life to ensure effective and persistent gut colonization.

While we found that arrival order influenced colonization outcomes for all tested community members, the strength of priority effects varied notably across strains. For instance, all strains of *Bifidobacterium asteroides*, *Bifidobacterium* sp. 2, *Lactobacillus helsingborgensis*, and *Lactobacillus kullabergensis* were almost entirely undetectable when introduced as part of the latecomer community. In contrast, strains from other species were still able to establish as latecomers, albeit at much lower abundances than when introduced first. The strain ES0263 of *Lactobacillus apis* stood out for its exceptional ability to successfully colonize even as a latecomer. The underlying reasons for these differences among strains remain to be elucidated, but may involve variation in niche overlap or the ability to interfere with established strains. Further investigation into the mechanisms behind these differences could reveal valuable traits for probiotic development or microbial engineering. These approaches are also of growing interest for promoting the health of managed honey bee populations (Motta et al., 2022).

A previous study demonstrated that phylogenetic relatedness predicts the extent of priority effects among microbes (Peay et al., 2011) i.e., priority effects were stronger between closer relatives, likely because they harbour greater niche overlap. Here, we tested this using dropout communities where one firstcomer strain was dropped at a time. Our expectation was that the latecomer strain of the same species would increase in conization if the priority effect was mediated through intra-specific competition. While this was the case for at least four of the six tested strains, we noted that the effect was relatively small (8–34%). In other words, the latecomer strain did not reach the same abundance as when arriving in the firstcomer community. Moreover, several strains of other species in the latecomer communities increased in abundance when a strain dropped out. These results suggest that priority effects are also mediated through interspecific interactions, in particular between species belonging to the same genus as the one dropped out. Hence, we surmise that not only strains of the same species but also those of different but closely related species have considerable niche overlap, and hence contribute to priority effects via niche pre-emption (Levy & Borenstein, 2013). Measuring the metabolic capabilities of these strains *in silico* and *in vitro* could be used to determine the niche overlap and subsequently predict community assembly and invasion. This was recently demonstrated in a study which showed that pathogens Klebsiella and Salmonella was less successful in invading communities containing particular species closely related to the pathogen in vitro and in vivo in the mouse gut (Spragge et al., 2023).

Our findings underscore the importance of arrival order in shaping gut microbial communities, particularly at the strain level and for the assembly of the gut microbiota of honeybees. While natural systems may buffer or modulate these effects through increased diversity and environmental complexity, the foundational role of early colonizers in determining community trajectories remains evident. These insights contribute to a broader understanding of how host-associated microbial communities are structured and maintained, with implications for both microbiome engineering in the context of bee health and ecological theory.

## ACKNOWLEDGMENTS

We thank the Lausanne Genomics Technology Facility (GTF) team at the University of Lausanne and the Next Generation Sequencing Platform of the University of Bern for performing the high-throughput sequencing for the experiments. We also thank Madalena Vaz Ferreira Real for providing helpful feedback on the manuscript. This work was supported by a Spirit grant (grant number 189496), an SNSF Consolidator grant (’GLOBEE,’ grant number 213860), and the Microbiomes, a National Centre of Competence in Research (grant number 180575 and 225148), funded by the Swiss National Science Foundation.

## AUTHOR CONTRIBUTIONS

Conceptualization – AP, GSM, FM, PE; Methodology – AP, GSM, FM, PE; Software – AP; Validation – AP; Formal analysis – AP; Investigation –AP, AS, GSM; Resources – PE; Data curation – AP; Writing - Original Draft – AP; Review and Editing –AP,GSM, FM, PE; Visualization – AP; Supervision – PE, FM; Project administration – PE; Funding acquisition – PE

## COMPETING INTERESTS

All authors declare they have no competing interests.

## DATA AVAILABILITY

Raw reads are deposited in the NCBI SRA database under the Project ID PRJNA1289778 (Reviewer link: https://dataview.ncbi.nlm.nih.gov/object/PRJNA1289778. Several intermediate files, including 16S Sequence database and RDS objects for analysis are included in the GitHub repository: https://github.com/Aiswarya-prasad/20240399_aprasad_PriorityEffects.

## CODE AVAILABILITY

All the code and intermediate files, including RDS objects, are found in the GitHub repository: https://github.com/Aiswarya-prasad/20240399_aprasad_PriorityEffects

## Notes

### Competing Interest Statement

The authors have declared no competing interest.

