## Supplementary Figures, Methods, Table legends for "Priority effects determine community composition at the strain level in the honeybee gut microbiota"

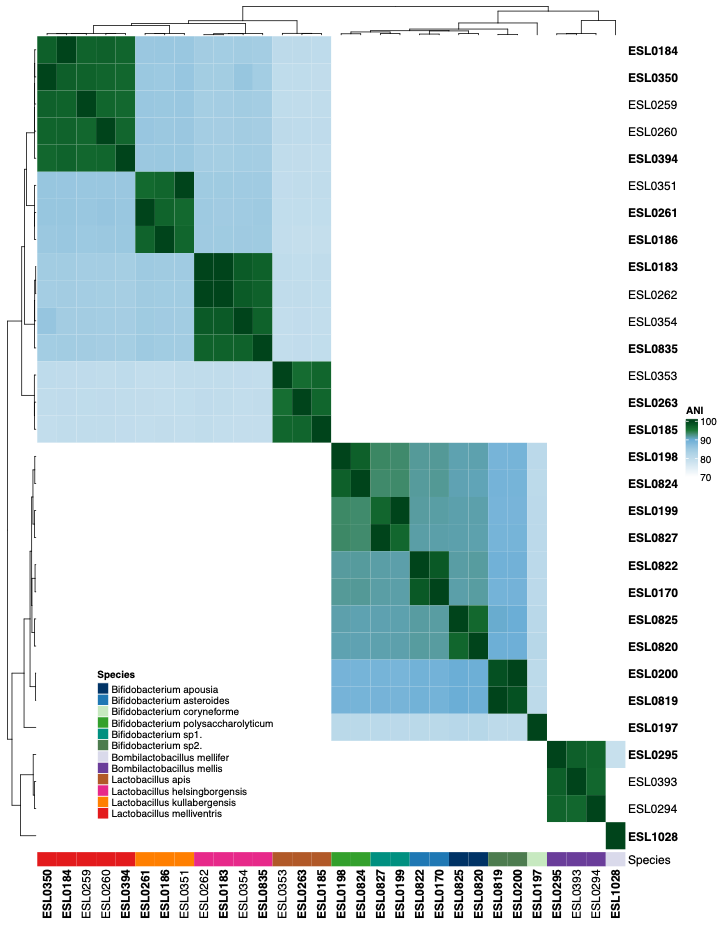

*Supplementary Figure S1. Heatmap of average nucleotide identity*. ANI heatmap of all strains clustered and annotated by species. Strains included in the experiments are marked by bold text.

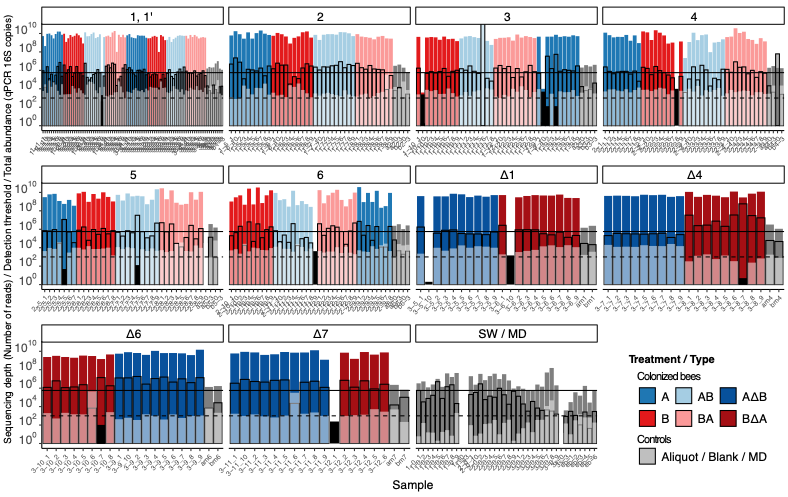

*Supplementary Figure S2. Total bacterial abundance, sequencing depth and detection limit across samples.* Total abundance of all strains per sample colored by treatment and facetted by the community combination. Colored bars with no outline represent the total bacterial abundance estimated using qPCR. White shaded bars without borders indicate the sampling depth in terms of the number of filtered reads. Solid black bars denote samples with low depth that were excluded from further analysis. The bars outlined in black represent the detection threshold for each sample established as the bacterial abundance corresponding to one sequencing read (i.e., the total bacterial 16S rRNA gene copies from qPCR divided by the total number of reads sequenced per sample). The solid horizontal line denotes the median detection threshold across all samples, and the dotted line the median depth per sample.

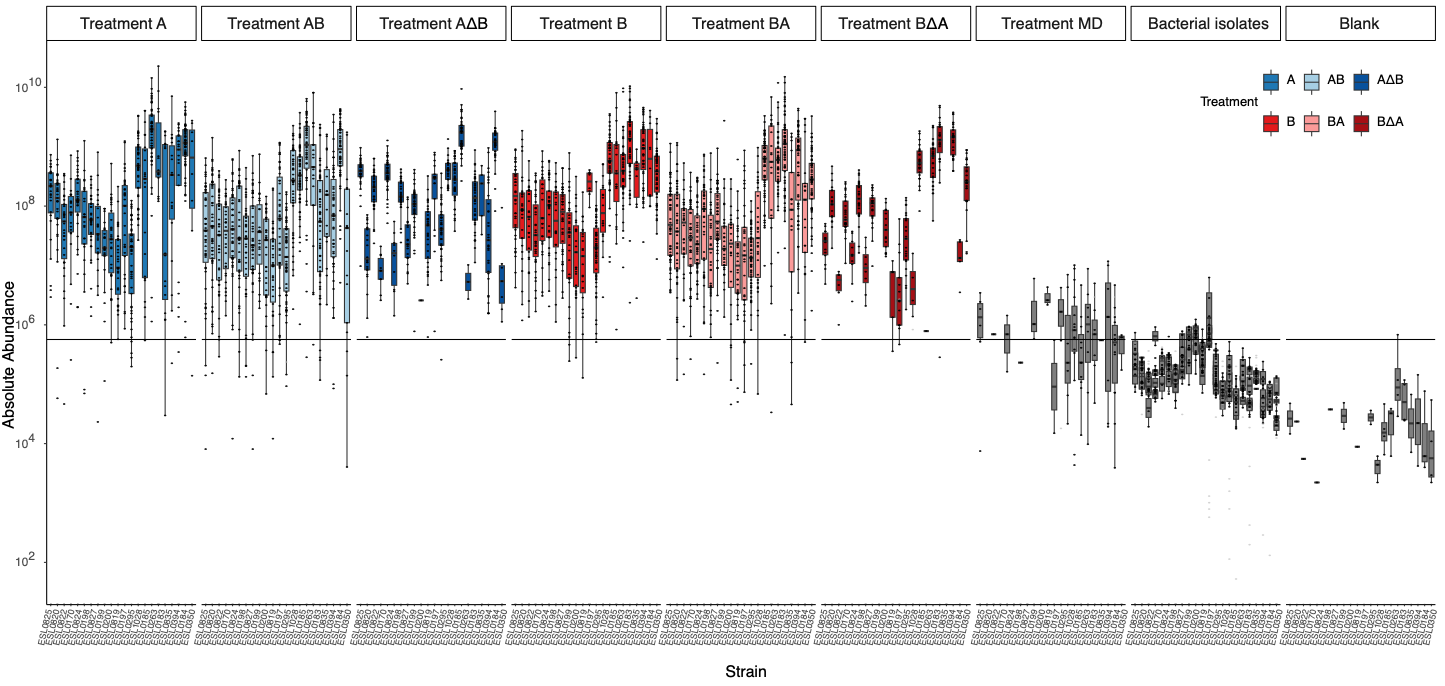

*Supplementary Figure S3. Abundance of strains in honeybee gut and control treatments.* Boxplots are based on the absolute abundance of each strain (on the x-axis) across different samples of each treatment, if it was above the detection threshold. Facets are made based on the treatment group for bee gut samples (MD treatment in grey), bacteria samples representing aliquots fed during the experiment, and blanks included during sample processing. Grey translucent points represent strains that were below the detection threshold in their respective samples. The horizontal line denotes the median detection threshold across all samples.

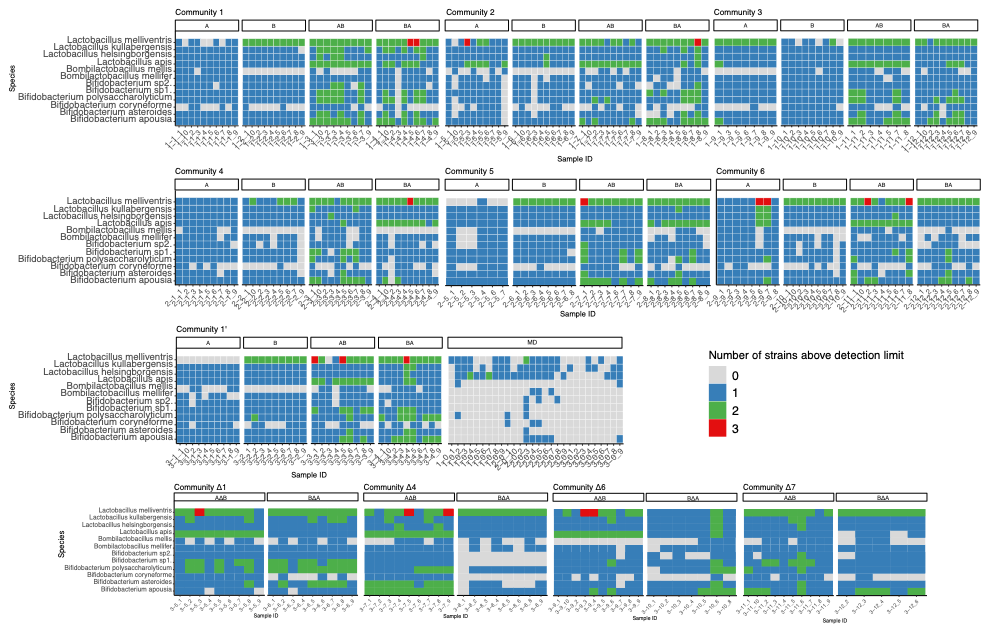

*Supplementary Figure S4. Heatmap of the number of strains detected in each sample.* Each row corresponds to a species, and each column represents a different honeybee sample. The heatmaps are faceted by experiment, community combination and treatments. The color on the heatmap indicates the number of strains of the species detected (absolute abundance above the detection threshold of the respective sample). In a majority of the bees, the facets A and B have one strain detected per species. In AB/BA, there are either 1 (strong priority effects) or 2 (with different abundances but above the detection limit for the respective samples). In some cases, 3 strains of *L. melliventris* are seen because one strain was misannotated as another species and hence included in some community combinations in addition to the two other strains of this species.

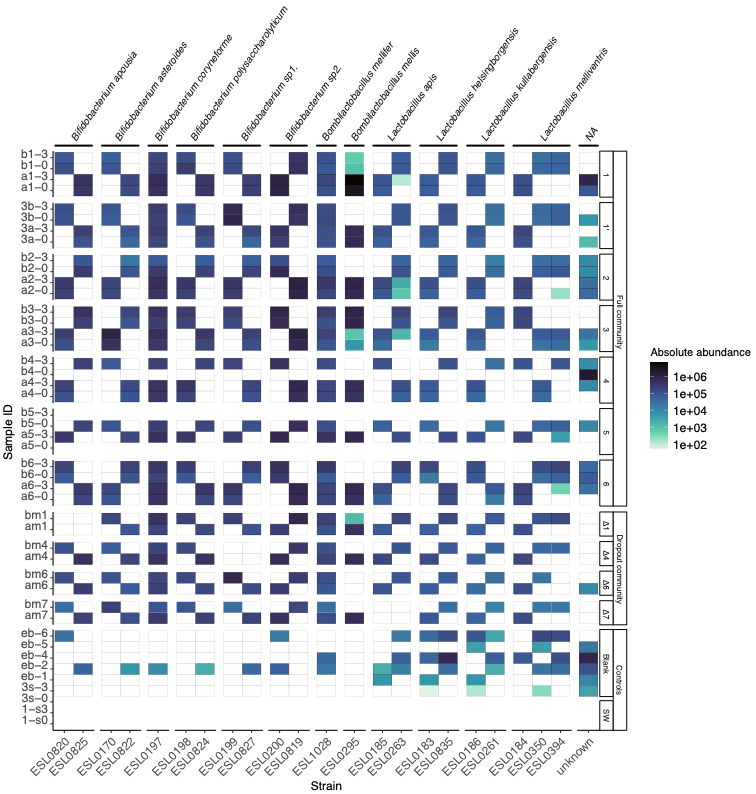

*Supplementary Figure S5. Heatmap showing strains detected in aliquots of inoculum.* Each column represents a strain, and each row is a different sample faceted by community type. For dropout communities, the aliquots for sequencing were collected from the tube used on Day 0, for the full community samples, an aliquot was taken from the tube used for inoculation on Day 0 and on Day 3 (sample IDs suffixed with ‘-0’ or ‘-3’). The controls comprised six extraction blanks (‘eb-’), which were samples of nuclease-free water added to the plates for DNA extraction and further processing. The others are samples of sterile sugar water from Day 0 and Day 3.

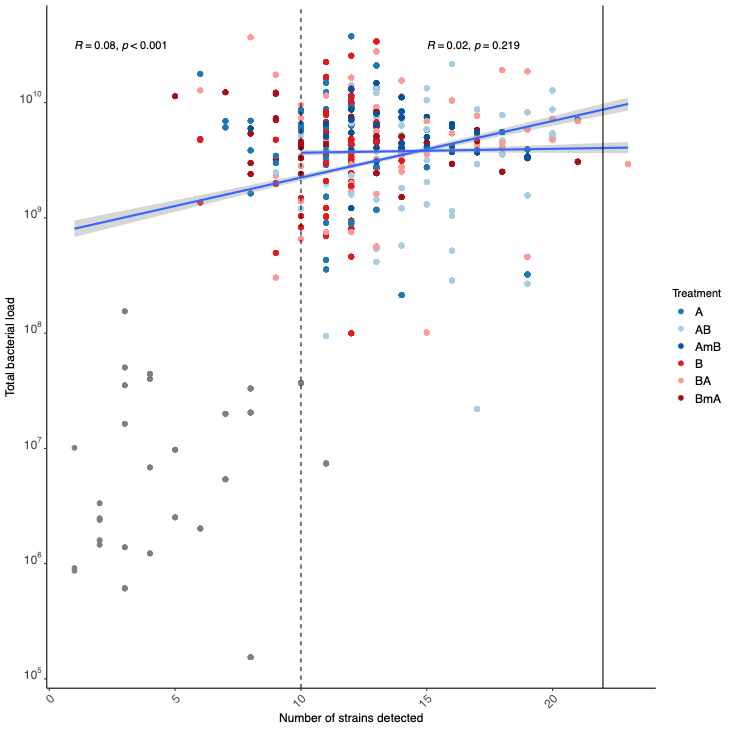

*Supplementary Figure S6. Correlation between the number of strains and bacterial load.* A scatter plot of the total 16S rRNA gene copies estimated by qPCR for each sample against the number of strains detected in the sample. The colors of the points indicate the treatment group to which the sample belongs. Grey points represent control treatments and are hence much lower in bacterial load. Spearman's ρ and p-value are estimated for including (left) and excluding (right) low bacterial load samples and those lacking of successful colonization of > 1 strain are displayed on the plot.

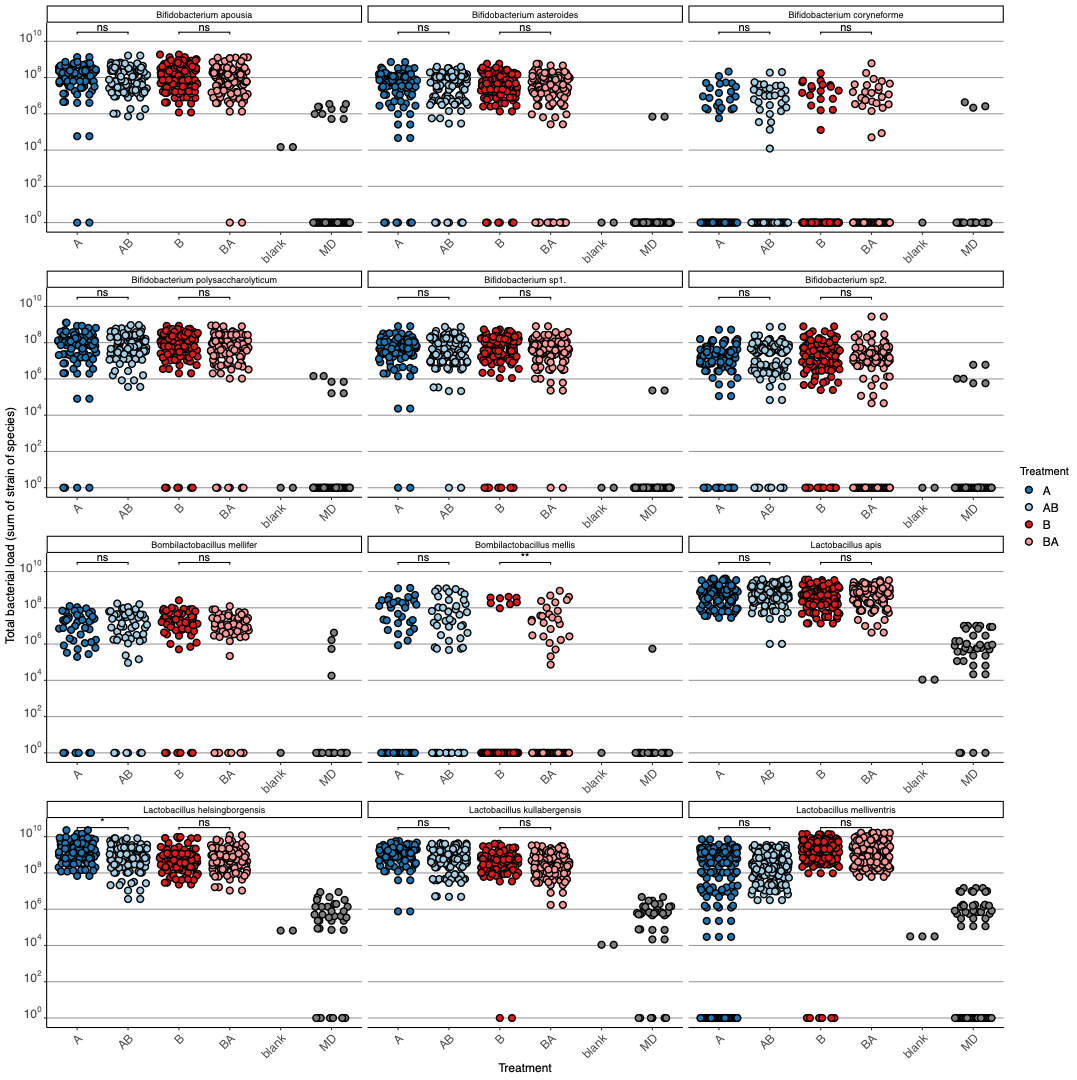

*Supplementary Figure S7. Total bacterial load of each species.* Absolute abundance of strains summed by species for each treatment group A and B, where only one strain per species was inoculated, and AB and BA, where two strains, one firstcomer (Day 0) and one latecomer (Day 3) were inoculated. Only points crossing the detection threshold for their respective sample are included. Horizontal lines represent the Benjamini-Hochberg adjusted p-value (p < 0.05) between treatments with one or two strains inoculated (A vs. AB and B vs. BA). Points where none of the strains of the species were detected were set to 1.

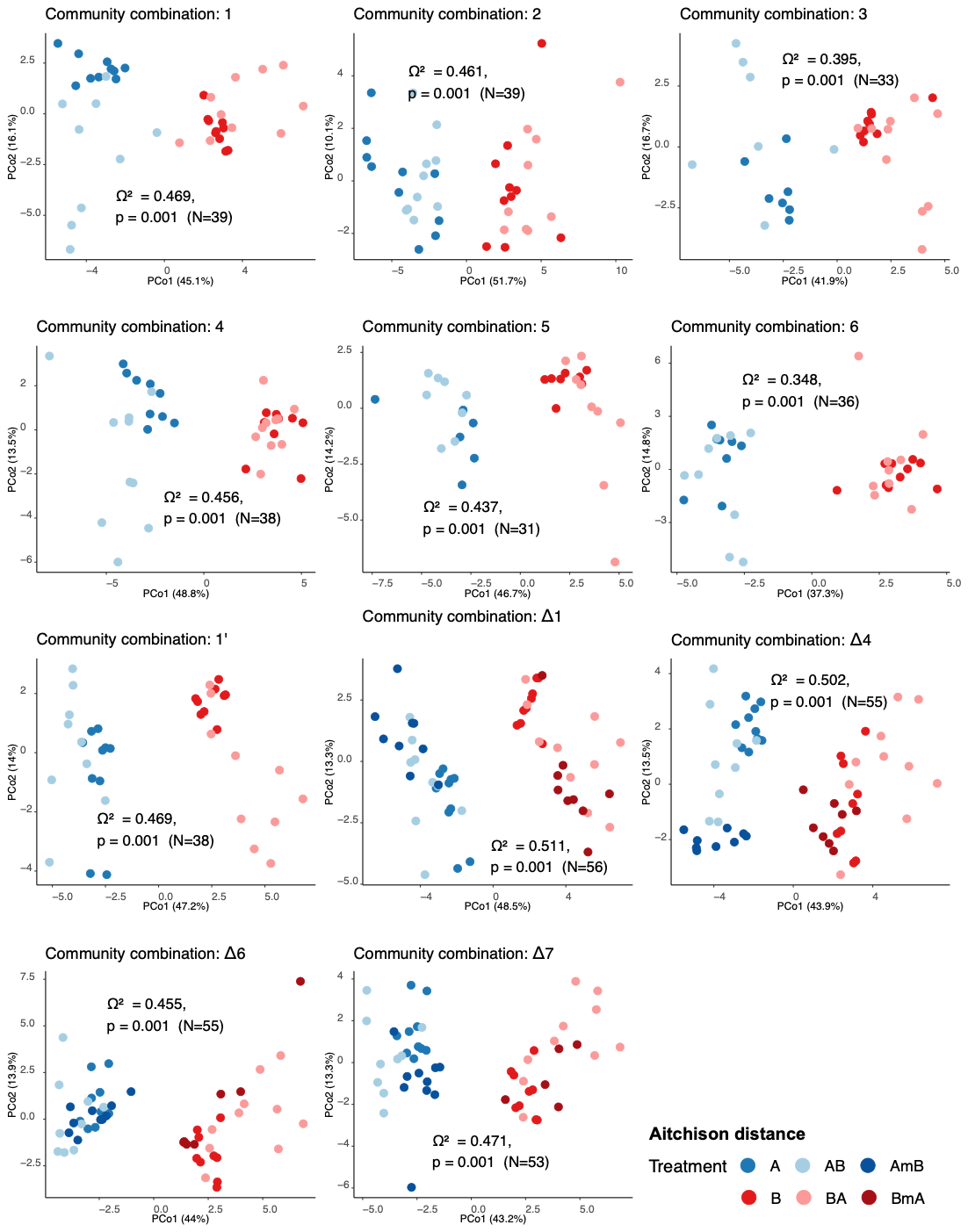

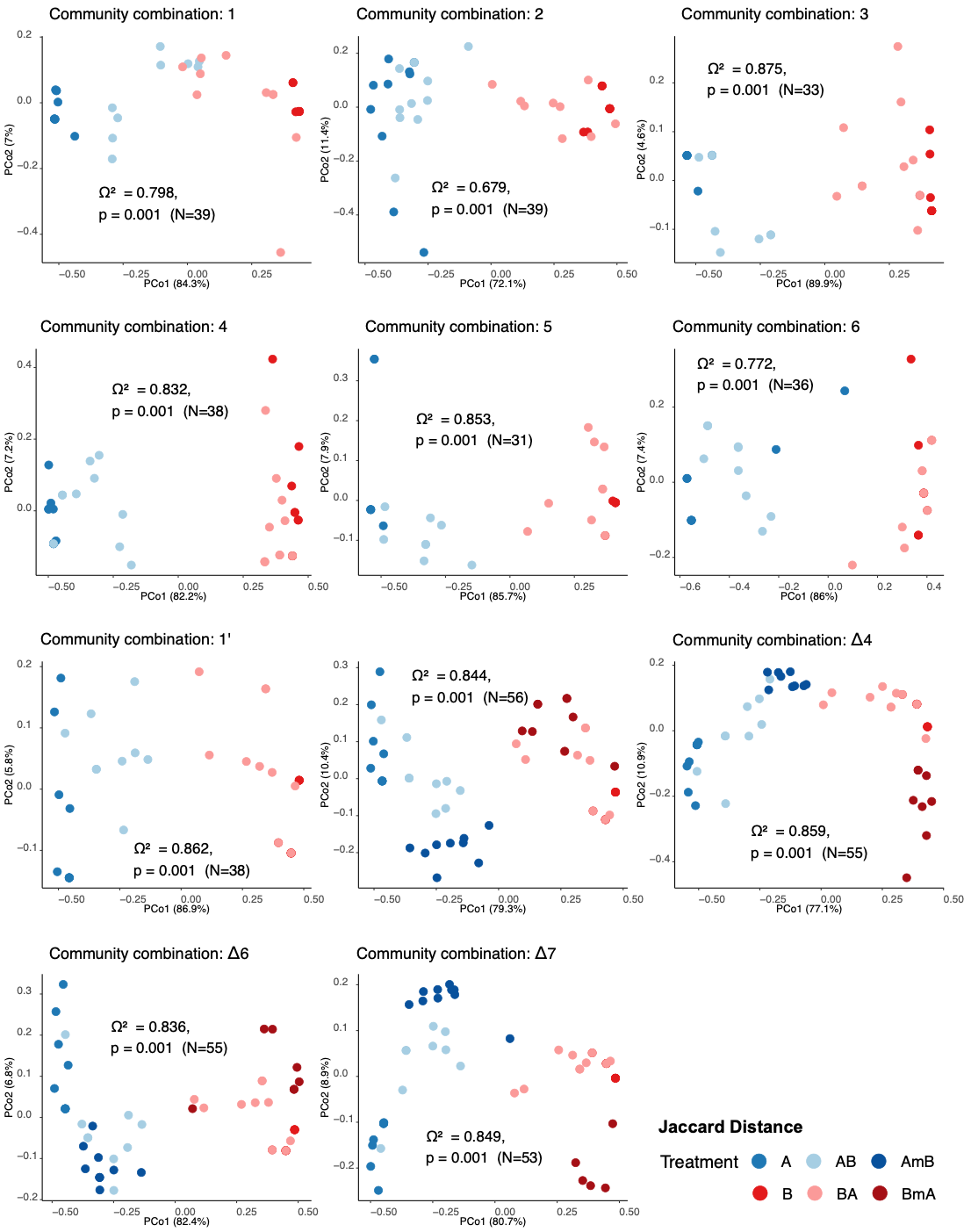

*Supplementary Figure S8. Community composition across samples.* PCoA plots to visualize the distance estimated by two different methods (Jaccard based on presence-absence and Aitchison based on relative abundance of strains) across samples inoculated with different community combinations across treatments A, B, AB, BA, and/or AΔ*B and BΔ*A.

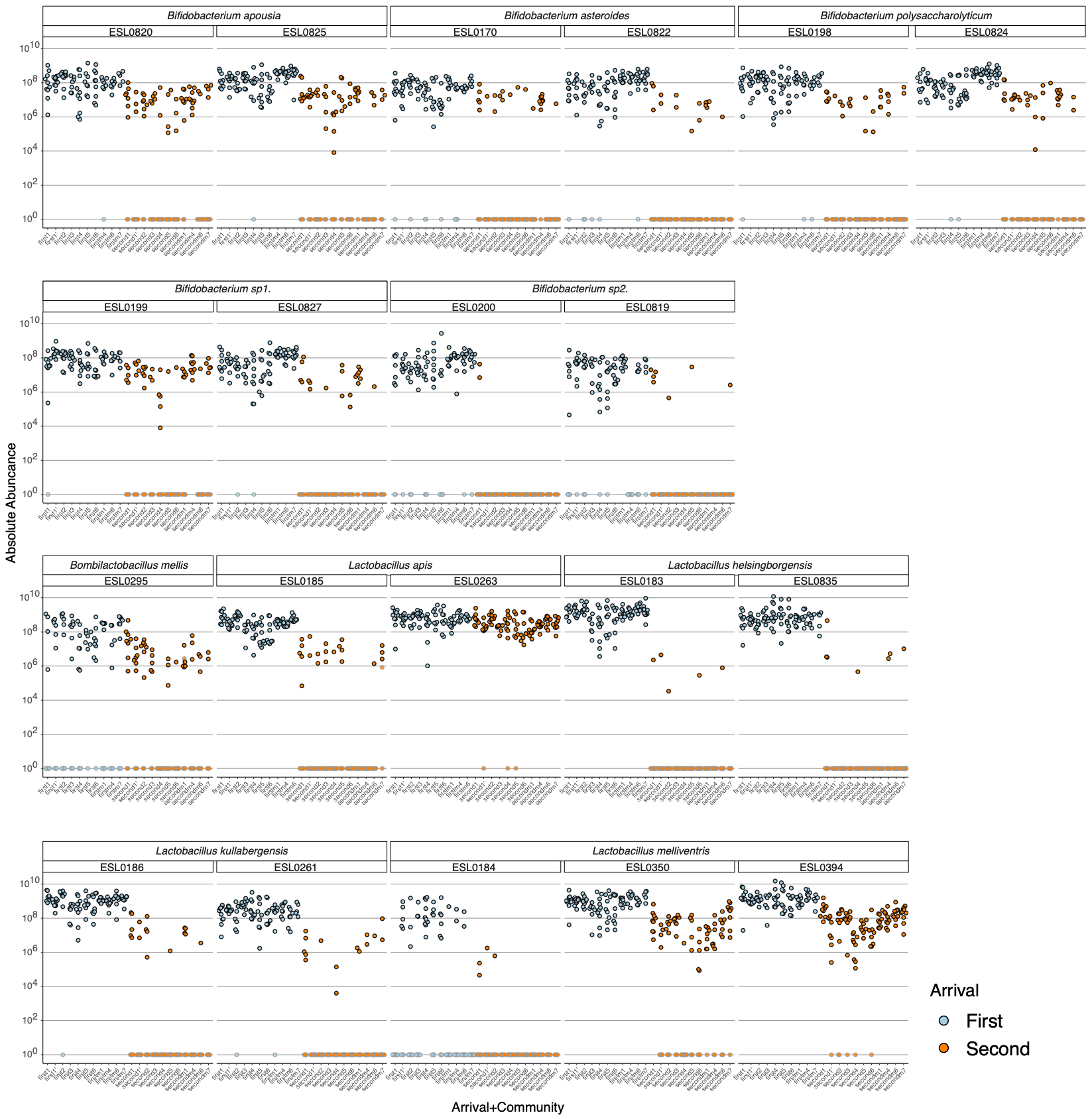

*Supplementary Figure S9. Colonization success of strains across treatments.* Absolute abundance of each strain across treatments where the strain was either in the firstcomer or latecomer community, also indicated by the color of the points. Points with grey outlines represent samples where the strain was not detected (below the detection threshold for that sample) and hence set to a value of 1.

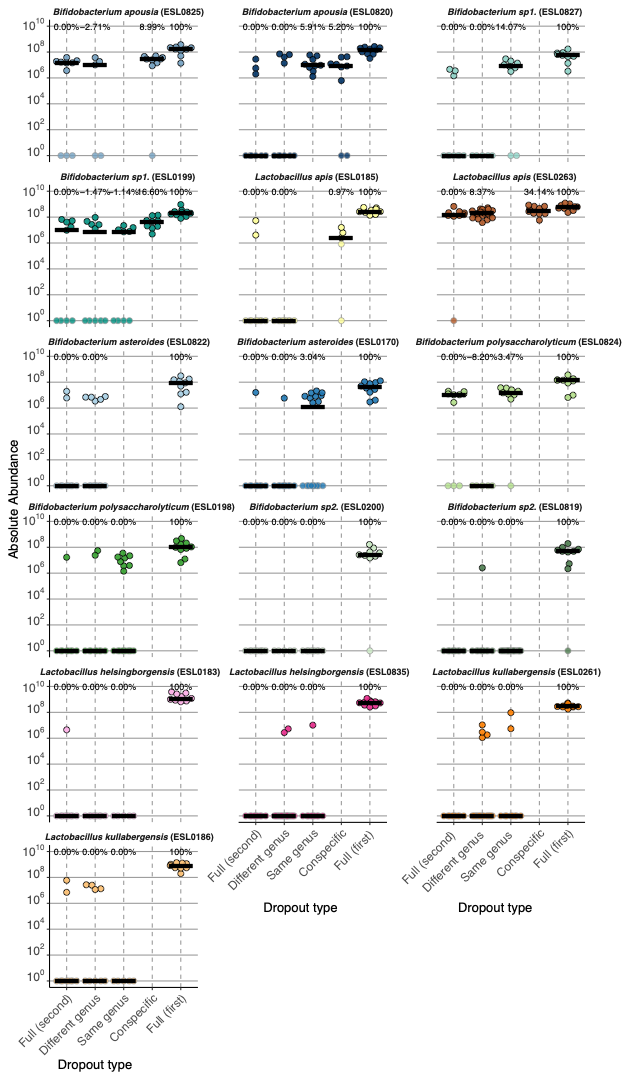

*Supplementary Figure S10. Effect of dropping out species from firstcomer community.* Abundance of each strain across full and dropout treatments. Shaded panels indicate strains that were present in community A and its dropout variations and hence, were firstcomers in A*B and second in B*A. Result of Wilcoxon rank sum test (two-sided) between abundance of latecomers in full and dropout treatments are annotated (* - p<0.05), complete results of statistical test including sample size per group are included separately (Supplementary Table S6). Small black arrows highlight dropout treatments where the latecomer showed better colonization success in terms of quantity or frequency. Points with grey outlines represent samples where the strain was not detected (below the detection threshold for that sample) and hence set to a value of 1.

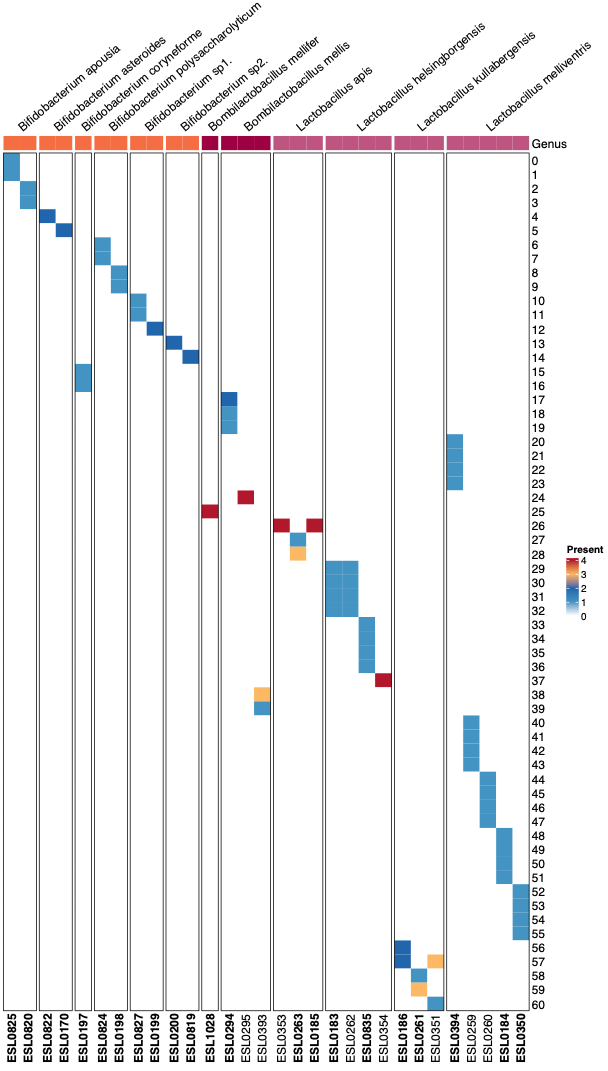

*Supplementary Figure S11. Heatmap of 16S rRNA gene sequence variants predicted across strains.* Each row represents a unique ASV of the 16S rRNA gene and each column a strain. The color of the boxes indicates the number of copies of that variant found in each respective strain as inferred based on Barrnap using the re-sequences long-read genome assemblies of these strains. Strains are faceted by the species they belong to and the bar on top colors panels based on the Genus. ASVs are named by the unique IDs (uid) assigned to them in our custom database. Strains included in the experiments are marked by bold text.

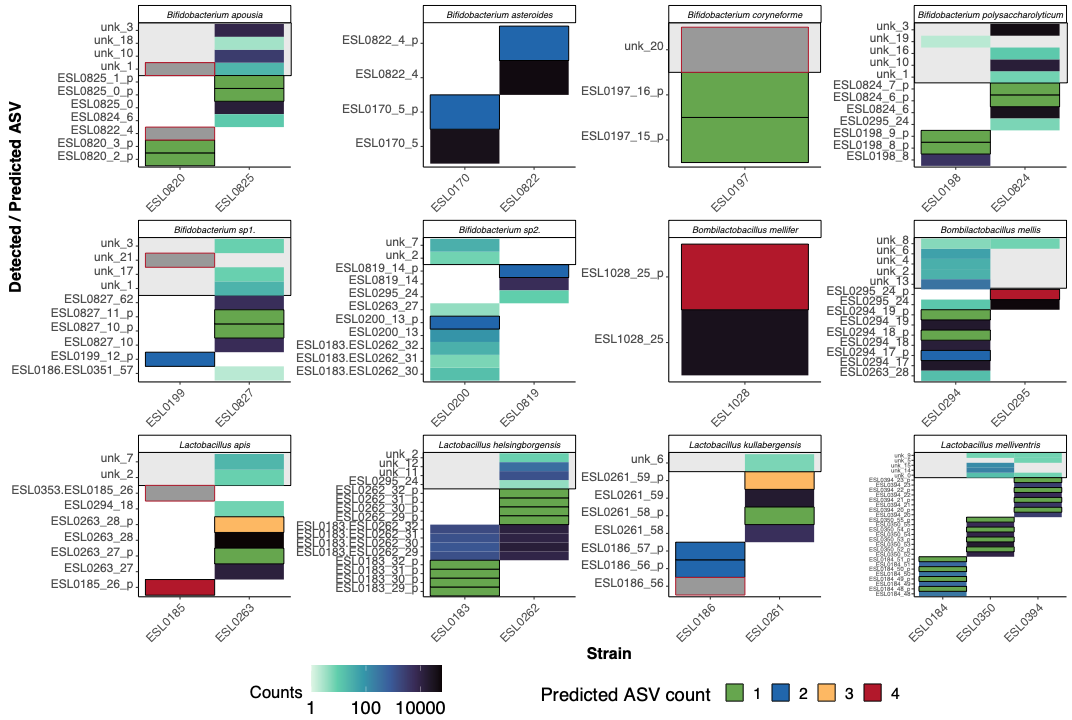

*Supplementary Figure S12 Predicted and detected ASV sequences and counts.* For each strain, copy number of predicted ASVs (marked by thick black boxes), and counts of detected sequences are shown faceted by species. Samples where the sequencing depth was too low for detection of ASVs (total reads <100) are indicated by a grey box (with red outline). On the vertical axis, ASV sequences predicted from the genome are suffixed with “_p”, their exact matches are named with the strain name and suffixed with their unique ID (_uid) in the custom database, and unknown sequences are prefixed with “unk_”.

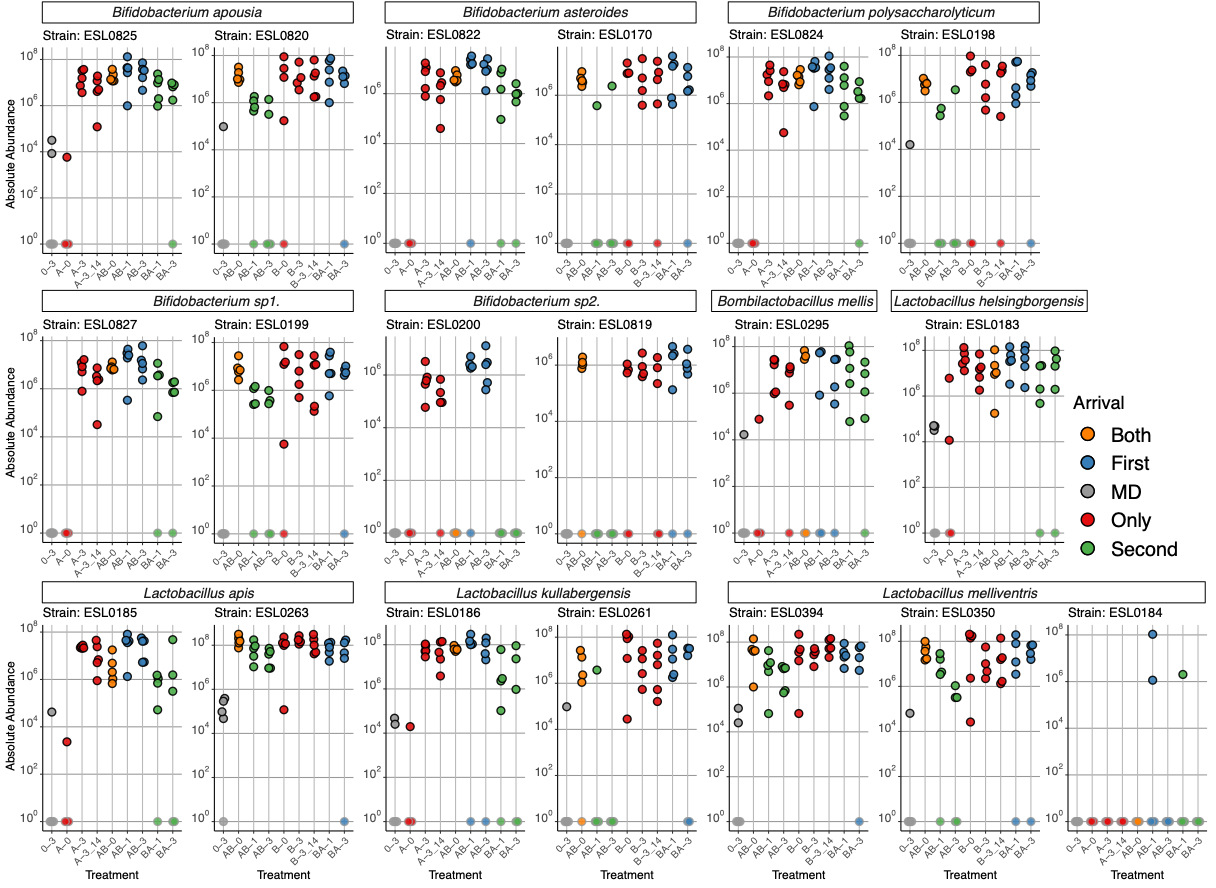

*Supplementary Figure S13. Pilot experiment.* Absolute abundance of each strain across treatments in the pilot experiment, colored and faceted by arrival order of the strain in the respective treatments. Points with grey outlines represent samples where the strain was not detected (below the detection threshold for that sample) and hence set to a value of 1.

### supplementary methods

#### strain selection and sequence database establishment

The bacterial strains used in this study were all isolated from the gut of adult bees of the Western honeybee (*Apis mellifera*) either in this or in previous studies (Brochet et al., 2021; Ellegaard et al., 2019). All strains belonged to one of twelve different species of the genera *Bifidobacterium*, *Lactobacillus,* and *Bombilactobacillus*, which are the predominant genera in the rectum of adult honeybees and typically co-occur in the same bee (Ellegaard & Engel, 2019; Maes et al., 2021; Prasad et al., 2025). More details about each strain can be found in **Supplementary Table S1**. We wanted to use the full-length 16S rRNA gene sequence to discriminate between these strains in the synthetic communities. However, the Illumina draft genomes available for some strains were inadequate for recovering complete 16S rRNA gene sequences, as divergent 16S rRNA gene copies were collapsed into a single contig during assembly. Hence, we re-sequenced most of the strains using long-read sequencing, either PacBio or Nanopore sequencing, and inferred 16S rRNA gene sequences and copy number from these genomes using Barrnap (**Supplementary Table S1, Supplementary Table S7**). We found that the V4 region (commonly used in previous studies) was identical for almost all our strains of the same species. However, outside this region, there were at least two distinct SNPs (sometimes >300 base pairs apart) in each 16S rRNA gene copy that could be used to differentiate most of the strains. The non-redundant full length 16S rRNA gene sequences of each genome were stored in a custom-made database and assigned unique IDs (uid, **Supplementary Table S7**). We then selected conspecific strains with the highest ANI that could still be distinguished by at least one 16S rRNA gene variant for our experiments (**Supplementary Fig. S10**, **Supplementary Fig. S11).**

From this strain collection, we assembled six synthetic communities, each comprising the same 12 species but differing in strain composition. For nine of the twelve species, we had two distinguishable strains available. However, for the two Bombilactobacillus species (*B. mellis,* and *B. mellifer*) and Bifidobacterium coryneforme, only one strain was available. So, these species were represented by the same strain in all communities, and as a consequence no priority effects could not be assessed for these species. However, we included them in the communities to avoid unoccupied niches that could be filled by others. Finally, the species *Lactobacillus melliventris* turned out to be represented by two strains (ESL0394 and ESL0350) in three of the six communities, due to a miss-annotation in our internal strain collection. For these reasons, priority effects could only be measured for 8 of the 12 species, i.e. those for which two strains were available and could be tested reciprocally in the AB and BA communities.

For the construction of the synthetic communities used in these experiments, we selected the closest relatives - which are expected to have the highest niche overlap and lowest fitness difference hence, most likely to see priority effects - within each species based on ANI (**Supplementary Fig. S10**), which were also distinguished by at least one unique 16S rRNA gene sequence from each other (**Supplementary Fig. S11**). ESL0262 and ESL0183 were determined to be indistinguishable (in the pilot experiment), as they contain the same four copies of the 16S rRNA gene, so only one of the strains was included in the main experiment. Finally, For the *Bombilactobacillus* species and for *Bifidobacterium coryneforme,* only one strain was included in all the experiments, and hence, the same strain was present in all the respective treatments across community combinations. This was done to avoid leaving niches available to others that these prevalent species might otherwise occupy. For *Lactobacillus melliventris,* one community of each pair included two strains, as one of its strains was mis-annotated as a *Bombilactobacillus mellis,* and in those communities, there was hence no representative of this species. Specifically, ESL0295 and ESL0394 were intended to be two representatives of *Bombilactobacillus mellis,* but re-sequencing and comparison of the genomes after the experiments were complete revealed that ESL0394 is a *Lactobacillus melliventris* genome. This mis-annotation in our internal database has been corrected, and all figures reflect the correct annotation. Hence, we exclude ESL0295 from the priority effect calculations since it was only present in one of each community pair. However, ESL0394 and ESL0350 were in the same community (except in combination 4), with ESL0184 being their conspecific counterpart. The dropout treatments AΔ6B and BΔ6A which were intended to be *Bombilactobacillus mellis* dropouts, were also excluded from comparisons.

#### validation of the custom database and strain detection approach

To validate our database (**Supplementary Table S7**) and strain detection approach, we PacBio sequenced the full-length 16S rRNA gene variants of each strain using purified genomic DNA of each strain and matched the most abundant amplicon sequence variants (ASVs) to our database. Other than a few strains with low sequencing depth (<100 reads), we detected the expected number of sequences in most samples (**Supplementary Fig. S12**). A few false positive ASVs were detected, but they were orders of magnitude less abundant than the ASVs of the expected sequences. To further confirm that they were false positives, we also compared the ASVs that did not match any strains to all the sequences inferred across all experiments (four different sequencing runs). For strain ESL0820, we detected two ASVs not present in the Nanopore assembly. Because these ASVs were consistently found across multiple samples containing ESL0820, whereas the assembly-derived variants were not, we updated our database to reflect the observed ASVs. Similarly, we added an ASV for ESL0827 to our database which was not found in the assembly.

We included two sequences *a posteriori* (uid 61 and 62) corresponding to the strain ESL0820 because it went undetected, as neither of the two sequences identified from its genome matched with any ASVs from the experiment. However, one ASV occurred exactly in the samples where ESL0820 was added and, upon further inspection, matched both uid 2 and 3 closely. It was not identical to either because it had both the unique SNPs of those two sequences. Since this ASV was consistently detected across several sequencing runs, we conclude that it is not an artifact (chimeric) but rather that there was an error in Nanopore genome assembly, resulting in two copies assembled, one with each SNP, instead of two identical copies, each containing both the SNPs. Hence, we added this sequence to the database as uid 61 expected to be present in two copies in ESL0820 instead of one copy each of uid 2 and 3. Similarly, among the two copies expected to be found in ESL0827, uid 10 was detected consistently, but never uid 11. Instead, another ASV sequence matching uid 11 closely, but for one SNP, was consistently detected. We added that ASV sequence to our database as uid 62, expected in one copy in ESL0827 instead of uid 11. Hence, we established and validated our database of distinct full-length 16S rRNA gene sequences from each of the strains used in this study.

#### pilot experment to validate experimental parameters

We conducted a pilot experiment with a community combination similar to the ones used in the main experiment (**Supplementary Table S2**) to inform our choice of experimental parameters. We included twelve treatments with one cage per treatment containing 10 honeybees, of which five were sequenced. To confirm that all our strains could establish themselves in the honeybee when inoculated simultaneously, we included Treatment AB-0, where both communities A and B were inoculated on Day 0 (**Supplementary Fig. S13**). All strains were able to establish themselves and grew to similar abundances. To verify how colonization is impaired by age, we colonized a set of bees with one community at day 0 and another set of bees with the same community at day 3 and observed comparable levels of bacterial colonization. This validated our assumption that reduced colonization success of the Day 3 community strains was due to the presence of bacteria belonging to the community fed on Day 0 (firstcomers) and not from any physiological trait of older honeybees (**Supplementary Fig. S13**). Treatments AB-1, AB-3, BA-1 and BA-3 demonstrated that the effect of challenging the firstcomers with latecomers at Day 1 or Day 3 was visible, with Day 3 showing a clearer difference for some strains (**Supplementary Fig. S13**). Hence, we chose to challenge bees with the latecomers on Day 3 in subsequent experiments. Finally, we also confirmed that the microbiota-deprived (MD) bees that were only inoculated with sterile sugar water and fed sugar water and pollen in their cage were largely free of gut microbes (**Supplementary Fig. S13**).

DNA extraction was carried out using the CTAB and Phenol-Chloroform-Isoamyl alcohol-based extraction method, as described before (Prasad et al., 2025). High molecular weight DNA used for long-read genome sequencing of isolates was obtained using the Maxwell® RSC instrument and Maxwell® RSC PureFood GMO and Authentication kit, using an additional bead-beating step in Qiagen Power Pro tubes for 10 minutes on a bench-top Vortex mixer.

#### quantification of total bacterial abundance and pcr yields using qpcr

All qPCR reactions were conducted in triplicate with a 10 μL total volume containing 0.2 μM of each forward and reverse primer, 1x SYBR® Select Master Mix (Applied Biosystems), and 1 μL of the sample. Thermal cycling conditions included a denaturation stage at 50°C for 2 minutes, followed by 95°C for 2 minutes, 40 cycles of 95°C for 15 seconds, and 60°C for 1 minute. Standard curves were prepared with serial dilutions of plasmid DNA as described previously (Kešnerová et al., 2017) to determine primer efficiency. Copies of bacterial targets in gut samples were hence calculated by applying these efficiencies to the qPCR results as $E^{i-C_{T}}$ ($E$ – primer efficiency calculated form the slope of the standard curve as ${10}^{\frac{-1}{slope}}$, $i$ – intercept from standard curve and $C_{T}$ – cycle threshold obtained from the qPCR run). $C_{T}$ values above 26.53 could not be distinguished from the blank in standard curves and were hence considered below the detection limit. Copy number was adjusted for the DNA extraction fraction volume, elution volume, and dilution factor to estimate the number of copies per bee using the code (qpcr_data_parse.R) provided in the associated GitHub repository.

16S rRNA PCR samples were quantified in the same way to quantify the PCR yield and dilute them to equimolar concentrations for PacBio library preparation.

#### details about amplicon sequencing analysis to determine strain abundance

Instead of arbitrarily picking one of the ASV counts and assuming it is the strain count or taking their mean or median we use an approach to infer consider the most likely strain count of X given the counts of its ASV(s). For example, if a strain has 2 unique ASVs in 2 copies a and b, then strain count X = a/2 = b/2. But due to technical issues a does not = b. In addition, say one of the ASVs is not just unique to strain X but also found in strain Y. So Y = a/2 = c = d (c and d is found in single copy in strain Y). So, framing the counts as linear equations helps get the most likely strain counts explained by the ASV counts addressing both the most likely values for X when a and b are not equal as well as handling cases where ASVs are not unique. This approach hence remains generalizable for more complex syncoms as well.

The observed ASV counts per sample (*A*) were modeled as a linear combination of strain copy profiles (*C*) and the unknown strain abundances (*S*), such that *A = C·S*. Because *C* is typically not square (there may be more than one unique ASV per strain), *S* was estimated by minimizing ||*CS*−*A*||² using the *np.linalg.lstsq* solver in Python. This was done for each sample such that A and S were the counts per ASV and counts per strain, respectively, for that sample. Matrix S from each sample was combined into the strain/species table used for further analysis. Reads from ASVs not matched to any sequences in the database were aggregated as unknown reads (6.25% or less of the total reads in 90% of the inoculated bee samples).

### supplementary tables

*Supplementary Table S1.* Information about each strain and its genomes. Associated ASVs provides a list of ASVs by Unique ID that are found in the respective genome. Repeated IDs in the list represent multiple copies of the same sequence in the genome. Sequences associated with ASV IDs are provided separately in Supplementary Table S7.

*Supplementary Table S2.* Composition of communities with examples of community combination pairs in treatments.

*Supplementary Table S3.* Result of Wilcoxon rank sum test (one-sided) carried out to compare latecomer strains across dropout and full community combinations in Experiment 3.

*Supplementary Table S4.* This table shows the absolute abundance of each strain across all individual bees within a given treatment. For example, the list for strain ESL0825 under the 'only' condition represents its abundance in all bees where community A (which includes ESL0825) was administered alone in community combination 1. The 'first' condition corresponds to its abundance in bees where community A was fed first in the AB sequential treatment. Values listed under '1′' are the reference abundances used for comparison with strains in the corresponding dropout treatments; these values are highlighted in bold.

*Supplementary Table S5.* Results of GLMM using the function `glmer` model log_10(AbsoluteAbundance) ~ ArrivalOrder + (1 | CommunityCombo), with a log link specified by `family = Gamma(link = "log")`

*Supplementary Table S6.* Result of wilcoxon rank sum test (two-sided) carried out to compare latecomer strains across full community combinations in In Figure 3.

*Supplementary Table S7.* Table of all copies of 16S full length Amplicon sequence either predicted from full genome sequences or obtained from sequencing of 16S rRNA amplicons of isolate strains.

*Supplementary Table S8.* Complete table of raw data per strain across all samples and treatments
